# Long-term experimental evolution of seasonal influenza A/H3N2 virus reveals diverse evolutionary potential

**DOI:** 10.64898/2026.10.26.761891

**Authors:** Annelies J. H. de Rooij, Katina D. Hulme, Sarah van Leeuwen, Karen de Haan, Laura E. van Groeningen, Marliek van Hoesel, Zandra C. Félix Garza, Dirk Eggink, Simon P. J. de Jong, Alvin X. Han, Colin A. Russell

**Author notes:** AJHR and KDH contributed equally. AXH and CAR contributed equally.

## Abstract

The repeatability of evolutionary outcomes is a fundamental question with implications for predicting the evolution pathogens like influenza viruses. Here, we tested whether seasonal A/H3N2 influenza virus evolution follows reproducible paths in a simplified setting where the natural cellular environment is maintained but adaptive immune selection is removed. We serially passaged replicate populations of an A/H3N2 virus for 1,200 generations across 12 independent differentiated human airway epithelium culture lines (three replicates of four donors) and performed deep sequencing to reconstruct the evolutionary dynamics. Rates of variant accumulation differed significantly among lines but not among donors, and parallel evolution was minimal. Populations generated polymorphisms not observed in the global human population, reversions to historical sequences, and changes at positions held near-invariant globally since 1968. These findings suggest A/H3N2 evolutionary trajectories are shaped less by intrinsic functional constraint than by selective context and help explain why predicting influenza virus evolution remains difficult.

## Introduction

Predicting seasonal influenza virus evolution remains a critical challenge in influenza virus control. Antigenic drift necessitates biannual vaccine updates, and current vaccine manufacturing timelines require strain selection 6-12 months ahead of actual virus circulation ^1,2^. Consequently, extensive work has characterised and monitored the evolutionary dynamics of influenza viruses and attempted to forecast their future course ^3–18^, focusing predominantly on the immunodominant surface glycoprotein haemagglutinin (HA) of A/H3N2 viruses circulating in humans. Yet despite seasonal influenza virus evolution in humans being a highly consequential system to understand, fundamental questions remain unresolved: how repeatable are the observed evolutionary patterns, how many evolutionary outcomes are possible at any given point in time, and can those outcomes be predicted?

The repeatability and predictability of evolutionary outcomes are fundamental questions in evolutionary biology^19^. Evolve and resequence (E&R) experiments, such as the long-term evolution experiment (LTEE) on *Escherichia coli*^*20*^, are designed to examine these central features of evolution, as well as assess the relationship between genotypic and phenotypic changes^21–23^. By removing or relaxing constraints that operate in nature, these experiments have revealed adaptive trajectories that remain unobserved or inaccessible in natural populations ^24^. While these studies have established general rules of microbial adaptation^25^, most are far removed from their natural evolutionary context. E&R approaches have been applied to influenza viruses to assess antigenic drift^26,27^, host adaptation^28–33^ and drug resistance ^34–38^; however, these studies were generally short-term and imposed strong directional selection, conditions that funnel evolution toward predictable adaptive endpoints rather than revealing the broader range of accessible evolutionary trajectories.

Here we examined the long-term evolutionary dynamics of seasonal A/H3N2 viruses in a simplified yet highly biologically relevant setting. We serially passaged the same A/H3N2 virus (i.e. A/Japan/WRAIR10159/2008) starting population in an *ex vivo* differentiated human airway epithelium (HAE) system using three replicates of four different donors (12 independent lines) for a total of ~1,200 generations. This system preserves the cellular environment of the primary site of influenza virus infection while excluding adaptive immunity, a dominant but heterogeneous selective constraint on A/H3N2 evolution in humans. Comparing the resulting trajectories with those observed in the globally circulating virus population allowed us to investigate the repeatability of evolution in a controlled setting when this constraint is removed, and to gauge how much of the potentially accessible evolutionary space decades of surveillance has actually sampled.

## Results

### Reconstruction of the observed molecular evolution shows diverse dynamics across experimental populations

We established 12 lines of *ex vivo* differentiated HAE cultures from 4 healthy donors and serially passaged the same A/H3N2 virus starting population for ~1,200 generations (i.e. 100 72-hour passages), while maintaining a 1,000 virion bottleneck at each passage to preserve viral diversity and a low infectious dose (Supplemental Figure S1). To monitor the virus’ molecular evolution, we sampled each population at every ~120^th^ generation (i.e. tenth passage) and identified *de novo* single nucleotide variants (SNVs) using whole genome sequencing and a custom bioinformatics pipeline (see methods). For each of the 121 samples (including the starting virus population), we sequenced two technical replicates, which showed high concordance between raw SNV frequencies (R^2^: 0.93 – 1). The true SNV frequency was established via maximum likelihood estimation of the binomial proportions across technical replicates. Median sequencing depth across all samples was 12,149× (inter quartile range (IQR): 6,480× − 21,830×).

As the 12 experimental populations evolved independently, we expected SNV diversity to vary within and across donors. Consistent with this, each experimental population acquired different SNVs across segments, with no consistent patterns within individual donors, across donors or across segments, revealing broad SNV diversity (Figure 1). Phylogenetic reconstruction of the population consensus sequences showed separate branches for each population, supporting the independent divergences (Extended Data Figure 1).

**Figure 1.**
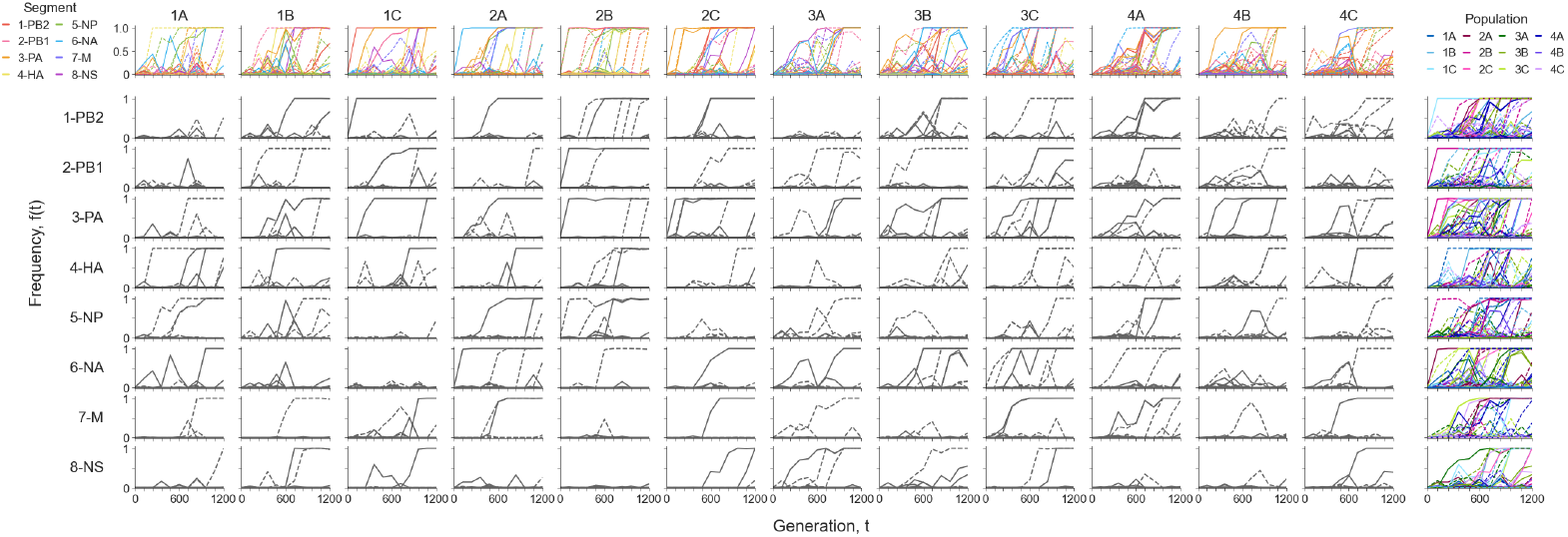
The allele frequency trajectories of all *de novo* single nucleotide variants (SNVs) detected in the experimental populations. Top row displays the allele frequency trajectories per population, with each trajectory coloured by the segment. Right most column displays the SNV trajectories per segment, with the trajectory coloured by the population. The remaining rows and columns display the trajectories per population per segment. Solid and dotted lines indicate non-synonymous and synonymous SNVs, respectively.

Since we do not know the appropriate population genetic model of the experimental populations, we employed a hidden Markov model (HMM; see methods) ^39^ as a phenomenological approach to estimate the approximate timing of emergence, fixation, and loss of SNVs. By the first sampling timepoint (generation ~120), both synonymous and non-synonymous SNVs had emerged in every population, though fixation had not yet occurred in all of them. The time to the first fixation event differed significantly by donor (analysis of deviance, *F* (3,8) = 5.88, *P=*0.02) (Extended Data Figure 2). Donors 1 and 2 averaged 240 and 200 generations, respectively, while donors 3 and 4 both averaged 560 generations. An exploratory contrast revealed that the first fixation event occurred 2.56 times later (geometric mean ratio) in donors 3 and 4 compared to donors 1 and 2 (*t* (8) = 4.25, Scheffé-adjusted *P* = 0.02, 95% confidence interval (95%-CI): 1.18 − 5.52). Individual pairwise differences were not significant after Holm correction.

Following the first fixation event, SNVs continued to fix over time, with the loss and fixation of SNVs matching the reductions in genetic diversity over time (Extended Data Figure 3). Notably, during the initial accumulation of genetic diversity (first 480 generations) the rate of diversity accumulation differed between donors (linear mixed model (LMM), overall interaction *F*(3) = 20.12, *P* = 2×10^−8^), with the donor 4 populations accumulating diversity faster than the other donor populations (Tukey-adjusted *P* < 0.005 for all three comparisons), and donor 3 populations accumulating diversity faster than those of donor 1(Tukey-adjusted *P* = 0.02); no other pairwise differences were significant.

To assess the rate of SNV accumulation over time, we calculated the total derived allele frequency^39^ for each population at each sampling timepoint (Figure 2A). Individual population rates exhibited significant variation in SNV accumulation (linear model overall interaction *F* (11) = 37.38; non-synonymous *F* (11) =14.97^;^ synonymous *F* (11) =44.11; all *P* < 2.2×10^−16^). Though these rates were comparable across donors (LMM, overall interaction *F* (3) = 0.72, *P* = 0.56), even when partitioned by substitution type (i.e., non-synonymous (*F* (3) 0.59, *P* = 0.64) vs synonymous (*F* (3) = 0.61, *P* = 0.62). When we further separated accumulation by genome segment, however, each population showed unique patterns, with no shared trends within donors or across segments (Figure 2B).

**Figure 2.**
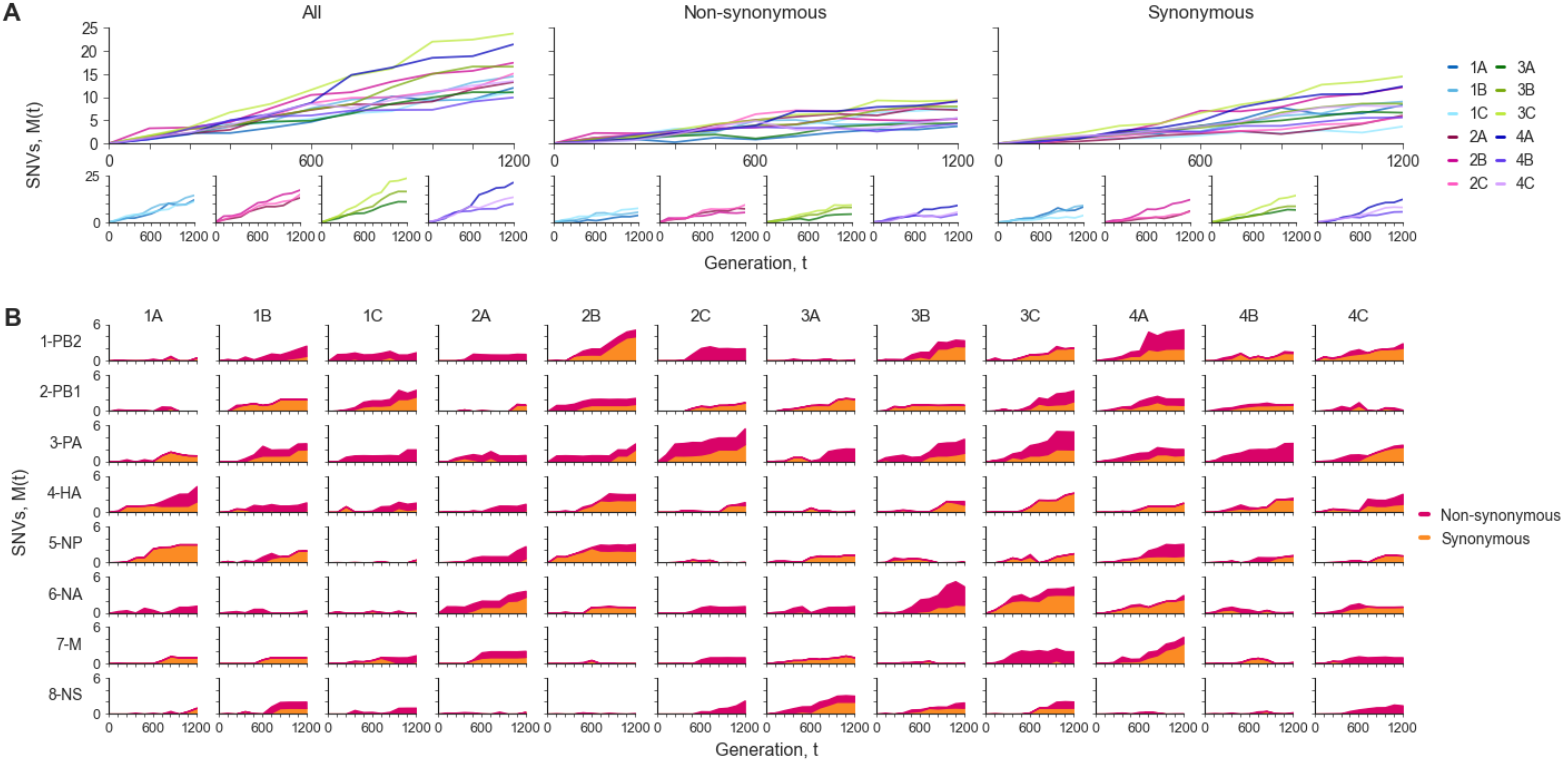
The accumulation of SNVs over time, measured by total derived allele frequency *M*_*p*_(*t*). **A**, Accumulation of all, non-synonymous, and synonymous SNVs across the genome per population *p*. Top panels display all populations, whereas the lower panels display the accumulation SNVs over time per donor. **B**, Accumulation of synonymous and non-synonymous SNVs over time per population per segment.

### Evidence against neutral evolution within all experimental populations

To test whether the evolutionary trajectories reflected genetic drift alone or were shaped by additional selective forces, we compared our experimental results to those of a drift-only forward-time simulation model of neutrally evolving populations that mirrored our experimental design. These simulations did not produce SNV fixation events (Figure 3A), suggesting that genetic drift alone is unlikely to account for the fixation events observed in our experimental populations. We used a statistical framework to classify SNVs from our experimental data as drift, transient, and fixed (Figure 3B); the transient and fixed trajectories deviated significantly from the simulation drift trajectories, providing evidence that the experimental populations are likely subjected to non-neutral evolutionary forces.

**Figure 3.**
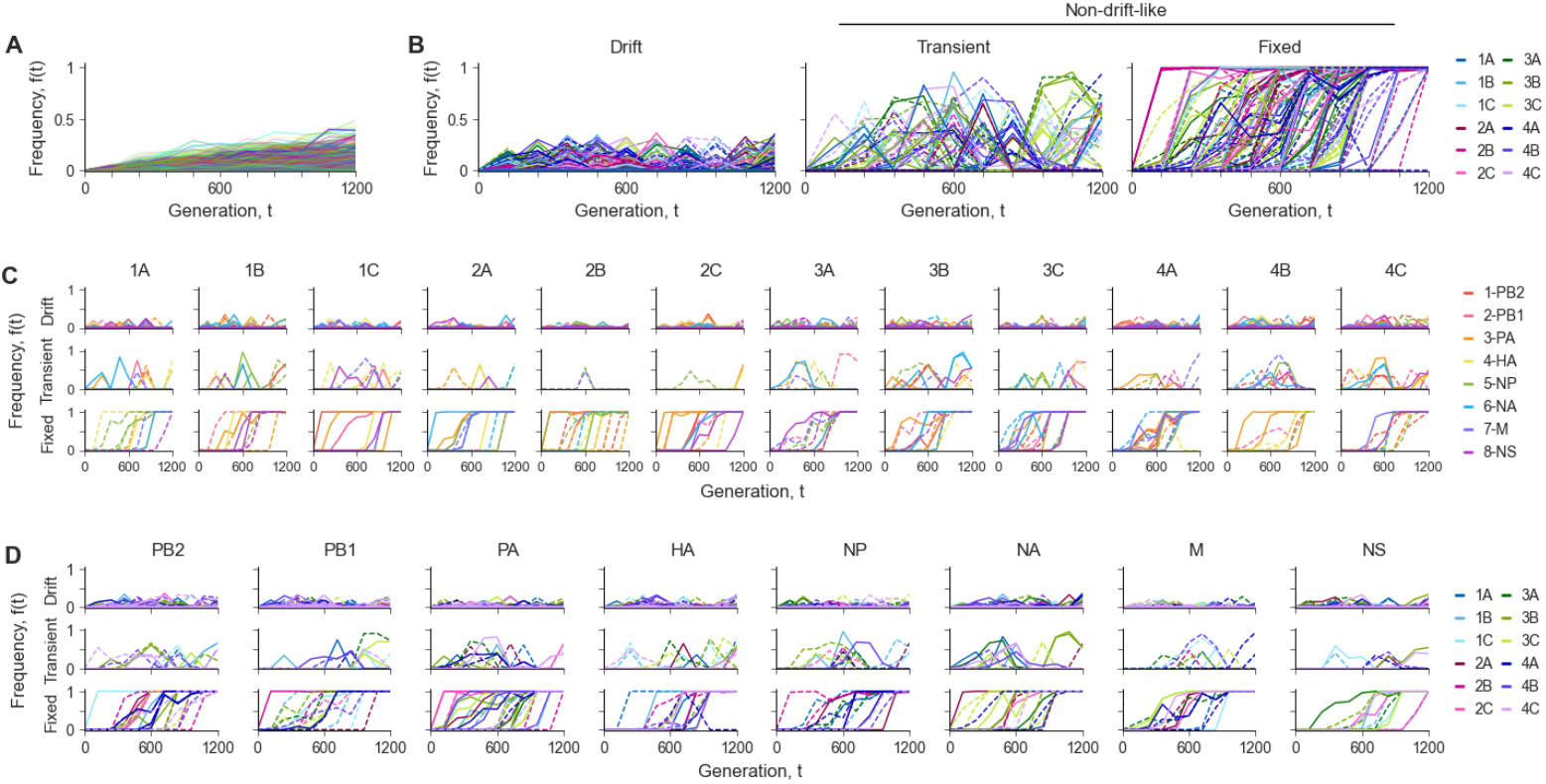
Frequency trajectories of observed SNVs classified based on the statistical inference of drift-like behaviour compared to simulated drift-like trajectories. **A**, simulated drift(-like) trajectories. **B**, Observed SNV frequency trajectories by SNV classification. From left to right: trajectories of SNVs with drift-like behaviour, SNVs with significant non-drift-like behaviour that did not fix, and (non-drift-like) fixed SNVs. SNV trajectories are coloured per population. **C**, SNV trajectories per population per SNV classification coloured by segment. **D**, SNV trajectories per segment per SNV classification coloured by population. Solid and dotted lines indicate non-synonymous and synonymous SNVs, respectively.

Non-neutral genetic signatures (i.e. transient or fixed trajectories) appeared in every population (Figure 3C), but varied widely by segment. Only five populations (1A, 1B, 2C, 3C, and 4C) carried non-drift-like SNVs (i.e. transient or fixed) across all segments, but none retained this pattern after filtering for non-synonymous substitutions (Figure 3C, Supplemental Table S3), indicating that non-drift-like changes were segment-specific rather than genome-wide. Non-synonymous non-drift-like SNVs showed no consistent pattern by donor (Figure 3D). Segments 3-PA, 1-PB2, and 6-NA were most commonly affected (12, 9, and 9 populations, respectively), typically with 1 to 4 SNVs each, suggesting broad tolerance for non-neutral change rather than parallel evolution. Conversely, 7-M had the fewest affected populations. Synonymous SNVs accumulated more uniformly across segments (Figure 3D, Supplemental Table S4), with the exception of 6-NA and 8-NS (both 7 populations only). For 8-NS, this is likely due to the 257 nucleotide overlap between the NEP and NS1 open reading frames (ORFs), which constrains synonymous changes.

### Non-neutral signatures persist after accounting for confounding factors

We then identified non-drift-like signatures that are likely confounded by to genetic linkage, mutation bias or codon usage bias (CUB). For linkage, we distinguished hitchhiking (i.e. a synonymous “passenger” SNV linked to a non-synonymous “driver” SNV) from co-occurrence (i.e. linkage between two SNVs of the same substitution type reflecting either hitchhiking or epistasis). Co-occurrence was reported without further interpretation, lacking the functional data to separate the mechanisms. Evidence for hitchhiking and/or co-occurrence was found in all populations (Figure 4A; annotated version in Supplemental Figure S2), but accounted only for 2-13 (10-39%) of the non-drift-like SNVs observed (Figure 4C).

**Figure 4.**
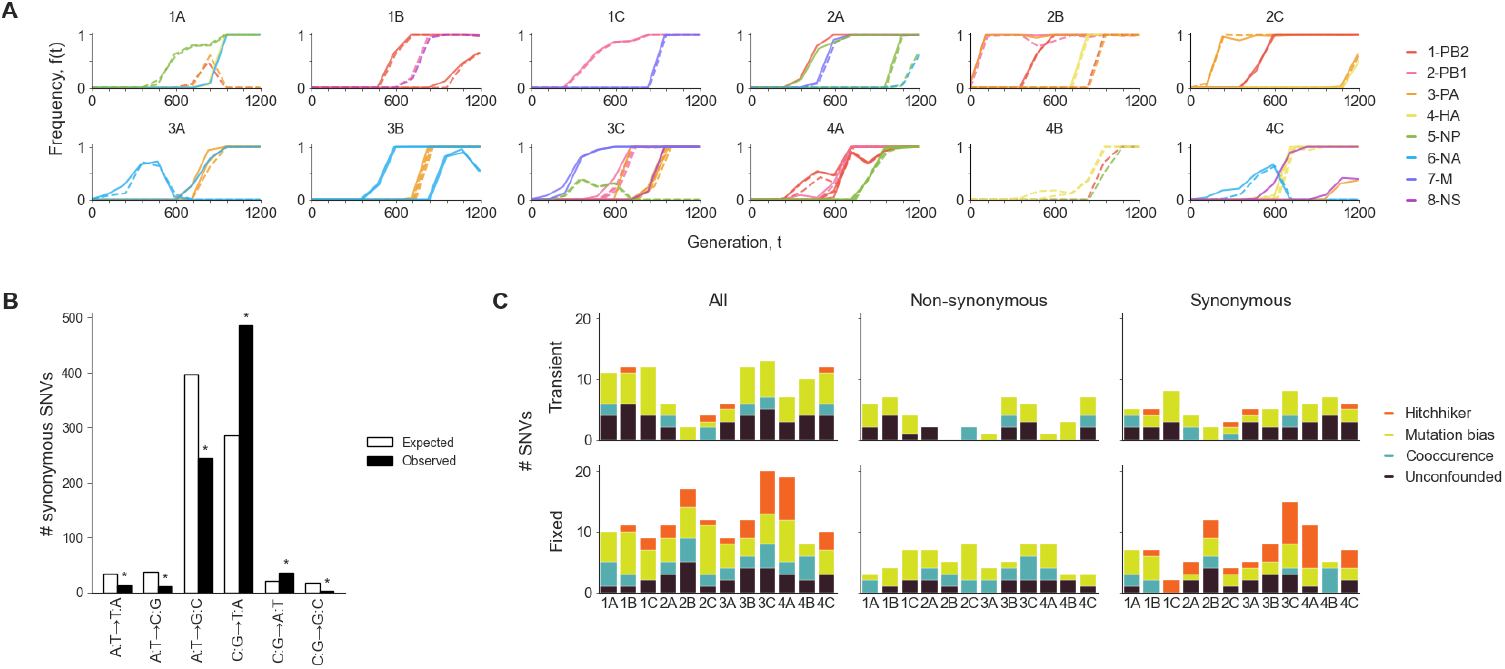
Decomposition of non-drift-like SNV trajectories by potential confounding factors. **A**, SNV frequency trajectories with evidence for putative linkage (both hitchhiker and cooccurring pairs) per population. SNV trajectories are coloured by segment. Solid and dotted lines indicate non-synonymous and synonymous SNVs, respectively. Annotated version in Supplemental Figure S2. **B**, Pooled expected and observed mutational spectrum for synonymous SNVs within the non-overlapping ORFs for PB1, PB2, PA, HA, NP, and NA. Asterisk indicates Holm-adjusted *P*<0.05 (binomial test). **C**, Number of SNVs by non-neutral classification and substitution type per population coloured by putative confounder. SNVs confounded by both putative linkage and mutation bias are coloured by linkage. Only the passenger in the hitchhiker pair is coloured, whereas both cooccurring SNVs within the pair are coloured.

Next, we assessed mutation bias by calculating the transition/transversion (Ti/Tv) ratio from all synonymous SNVs. As all populations had an elevated Ti/Tv ratio relative to the expected ~3 ^40,41^ (binomial tests, *P*<0.05 for all except 2B; Supplemental Table S5), we estimated an experiment-wide Ti/Tv ratio using a binomial generalised LMM (GLMM) at 11.76 (95%-CI: 9.15 – 15.48), which was also significantly elevated from the expected ratio (likelihood ratio test (LRT), *χ*^2^ (1) = 31.59, *P* =1.9×10^−8^). Next, we tested which mutation types drove this bias. Mutational spectrum counts were comparable across populations (*χ*^2^ = 41.63, Monte Carlo *P* = 0.9152), and therefore pooled (*n* = 791). For transitions, C:G→T:A mutations were significantly elevated (two-tailed binomial test, Holm-adjusted *P* = 9×10^−46^), while A:T→G:C were significantly reduced (two-tailed binomial test, Holm-adjusted *P* = 7.8×10^−27^). Among transversions, C:G→A:T was significantly elevated (two-tailed binomial test, Holm-adjusted *P* = 0.0016). The increased bias towards C:G→T:A mutations held across populations (Extended Data Figure 4), substitution types, and classifications (Extended Data Figure 5), so we labelled C:G→T:A SNVs as possible mutation bias, capturing 9-19 SNVs (47-88%) per populations, of which between 1-8 SNVs overlapped with identified linkage events.

Finally, we found no evidence that CUB explained the fate of the non-drift-like synonymous SNVs or the skewed Ti/Tv ratio’s (Extended Data Figure 6). After accounting for all potential confounders, 1-9 SNVs (6-33%) per population still remained unexplained (Figure 4C), suggesting persistent signatures of likely non-neutral evolution.

### Haplotype reconstruction reveals distinct dynamics within and between experimental populations

To further assess the evolutionary dynamics within and between experimental populations, we reconstructed the haplotypes by genome segment over time (Figure 5). Haplotype reconstruction extends beyond single-locus frequency trajectories, enabling inference of genome segment-level dynamics. For this analysis, we included all non-drift-like SNVs when their frequency was above 5%; since confounding factors could only be confirmed with functional data, we retained all non-drift-like SNVs to capture the full range of selection dynamics.

**Figure 5.**
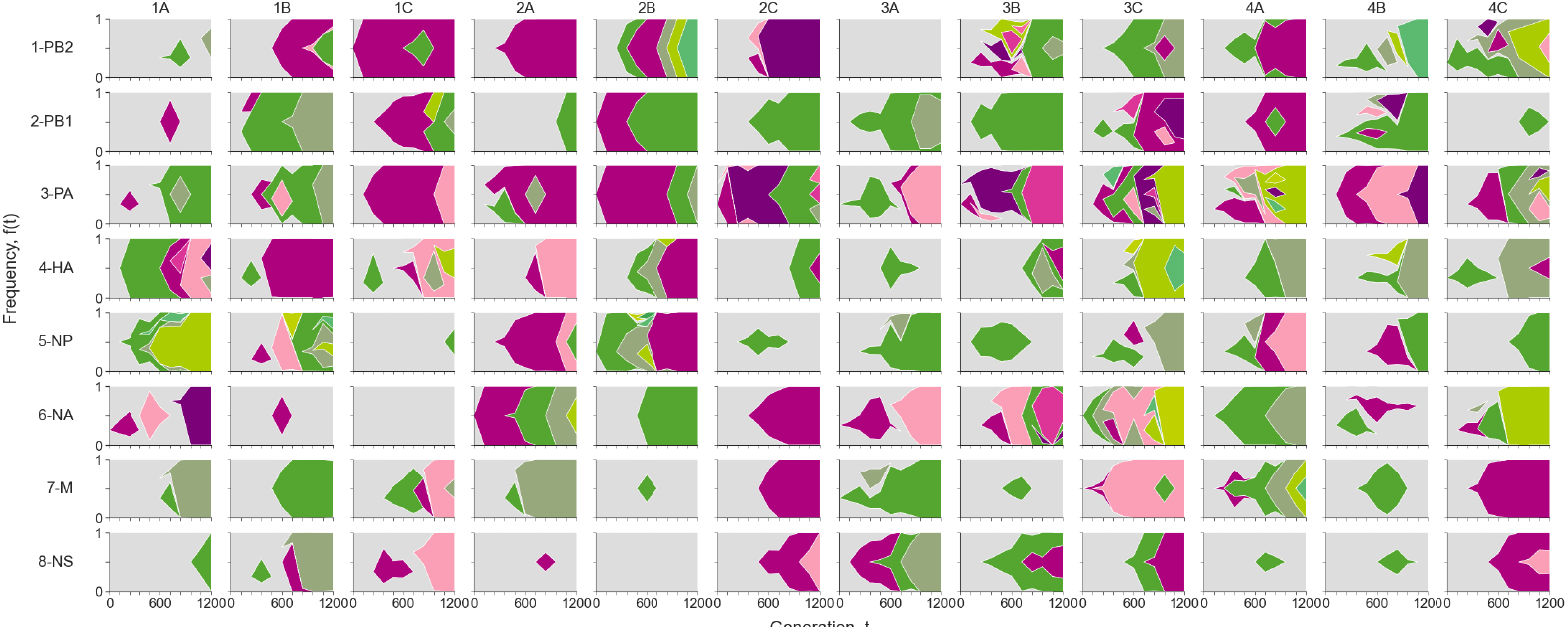
Dynamics of reconstructed haplotypes of non-drift-like SNVs within each population per segment. Within each subplot, each haplotype is shown in a unique shade of either pink or green. The colour of the haplotype denotes whether the latest SNV gained within the haplotype was non-synonymous (pink) or synonymous (green). In case of multiple gained SNVs within a haplotype, the haplotype is coloured pink if one of these SNVs was non-synonymous; otherwise, it is coloured green. Only haplotype observations above 5% frequency and only haplotypes with at least one observation above 10% frequency are plotted. The grey background shade indicates the ancestral haplotype.

We observed diverse dynamics across population and segments, including selective sweeps, clonal interference, hitchhiking, and/or epistasis (Figure 5). Segments within a population typically evolved independently, though concurrent changes occasionally occurred, such as an SNV loss in one segment coinciding with a sweep in another (e.g., population 3C, segments 4-HA and 5-NP), or simultaneous sweeps in different segments (e.g., population 2B, segments 2-PB1 and 3-PA). Sweeps were generally confined to one or two segments rather than spreading genome-wide. Simultaneous sweeps usually matched identified linkage events (Figure 4A). In some cases, however, sweeps cooccurred without identifiable linkage (e.g., population 1C, segments 7-M and 8-NS); here, individual SNV trajectories could not confirm linkage, though the cooccurrence itself still suggests hitchhiking or epistasis. We also found evidence of clonal interference, where multiple haplotypes arose concurrently (usually within the same genetic background) but only one eventually fixed (e.g., population 3C, segment 3-PA). In several populations, haplotypes persisted at low frequencies before acquiring additional SNVs, a pattern suggestive of either hitchhiking or epistatic effects and supporting interactions beyond those already identified through linkage. Consistent with the continuous fixation events observed across populations, no populations showed stably coexisting lineages. Together, these dynamics indicate that the evolutionary landscape was shaped primarily by independent evolution of individual segments rather than genome-wide selection.

### Minimal evidence for parallel evolution

Next, we investigated whether the populations had common paths of evolution. Most of the long-term E&R studies performed to date have examined organisms with much larger genomes and gene sets than influenza A/H3N2 viruses ^39,42^. These studies typically infer signatures of parallel evolution by identifying genes with more point mutations (and sometimes indels) than expected by chance ^39,42,43^. However, this approach is poorly suited to influenza A/H3N2 viruses, given its small genome, few genes, and high mutation rate. Instead, we tested whether the number of loci within a gene mutated in at least two populations exceeded neutral expectation, using a permutation-based null.

We evaluated parallelism across the non-overlapping ORFs of the PB2, PB1, PA, HA, NP and NA genes, focussing initially on non-drift-like non-synonymous SNVs. Only PB2 showed significant excess parallelism under the multiplicity-weighted statistic (*P* = 0.0010; Extended Data Table 1). This signal was driven by a single SNV (G2218A; D740N) exhibiting non-drift-like signatures in three populations (2C, 3B and 4A; Supplemental Table S6) and neutral drift in two others but was not significant under the site-count statistic (*P* = 0.056).

However, D740N is also flagged as a high-confidence co-occurring mutation in population 2C, indicating linkage, leaving only 3B and 4A as unambiguous unlinked signatures, which suggests that the support for parallelism is weaker than the multiplicity-weighted statistic implies. HA showed a single non-synonymous parallel SNV (G46A; A16T (signal peptide), populations 1B and 1C), but this was not significant for either statistic (*P* = 0.0624). No other gene showed evidence for parallelism.

For completeness, we also examined the overlapping ORF pairs M1/M2 and NS1/NEP, that were excluded from the primary analysis since SNVs in shared coding sequence cannot be unambiguously assigned to one gene. M1 showed significant parallelism (*P* = 0.0183 both statistics; Extended Data Table 1), driven by a SNV outside of the M2 ORF (C653T; A218V; population 2A and 4C; Supplemental Table S6), making this signal specific to M1. NEP also showed significant parallelism (*P* = 0.0107 both statistics), driven by a single SNV (G611A; E52K; population 2C and 4C) within the overlap with NS1 (R204K). NEP only accumulated one single-site SNV, while NS1 accumulated 8 single-site SNVs and showed a trend toward parallelism (*P* = 0.0771 both statistics). As position 611 is shared between both ORFs, this parallelism could not be attributed specifically to NEP or NS1.

Since synonymous SNVs are generally assumed neutral, we also tested them for parallelism; significance here could indicate site specific mutation bias or selection (e.g., on RNA secondary structure). Although some genes shared non-drift-like synonymous SNVs (Supplemental Table S7), we found no significant evidence for parallelism under either statistic (Extended Data Table 2).

Overall, evidence for genetic parallelism across populations was minimal, limited to a non-synonymous site in PB2, a non-synonymous site in M1, and a non-synonymous site in the overlap of NEP and NS1. Notably, all but one of the shared SNVs occurred across multiple donors, indicating that convergence was not donor-specific.

The broader lack of parallelism suggests non-neutral evolution did not follow a common trajectory across experimental populations or that the signatures of parallel evolution in our experimental system are too weak to be detected with our approach.

### Experimental populations explore trajectories distinct from those observed globally

Next, we compared the evolutionary pattern observed in our experimental system to historical global A/H3N2 virus evolution, focusing on non-drift-like amino acid (AA) mutations (i.e. non-synonymous SNVs). In all experimental populations, AA mutations occurred at globally conserved positions (i.e. single AA at ≥90% frequency at each point in time) (Figure 6A). By gene, most unique AA mutations occurred at globally conserved positions with exception of HA (4/13 AA mutations at globally conserved positions), M2 (1/3), and NS1 (4/9) (Supplemental Table S8 and Table S9). Notably, several of these changes were non-conservative (based on negative BLOSUM62 scoring) globally (16/66, 8 fixed; Supplemental Table S9). The emergence of non-drift-like AA changes in our experimental population at otherwise conserved positions suggests that alternative evolutionary trajectories are accessible, potentially optimising (non-antigenic) phenotypes not observed or inaccessible in natural populations. Additionally, in nine populations AA mutations occurred at non-conserved sites, but where the specific AA variant observed never reached > 10% global frequency (Figure 6A; Extended Data Figure 7). These changes were mostly conservative, but multiple non-conservative changes occurred in HA, and one each in PA, NA, and M2, further supporting divergence from observed evolutionary trajectories in the global human population.

**Figure 6.**
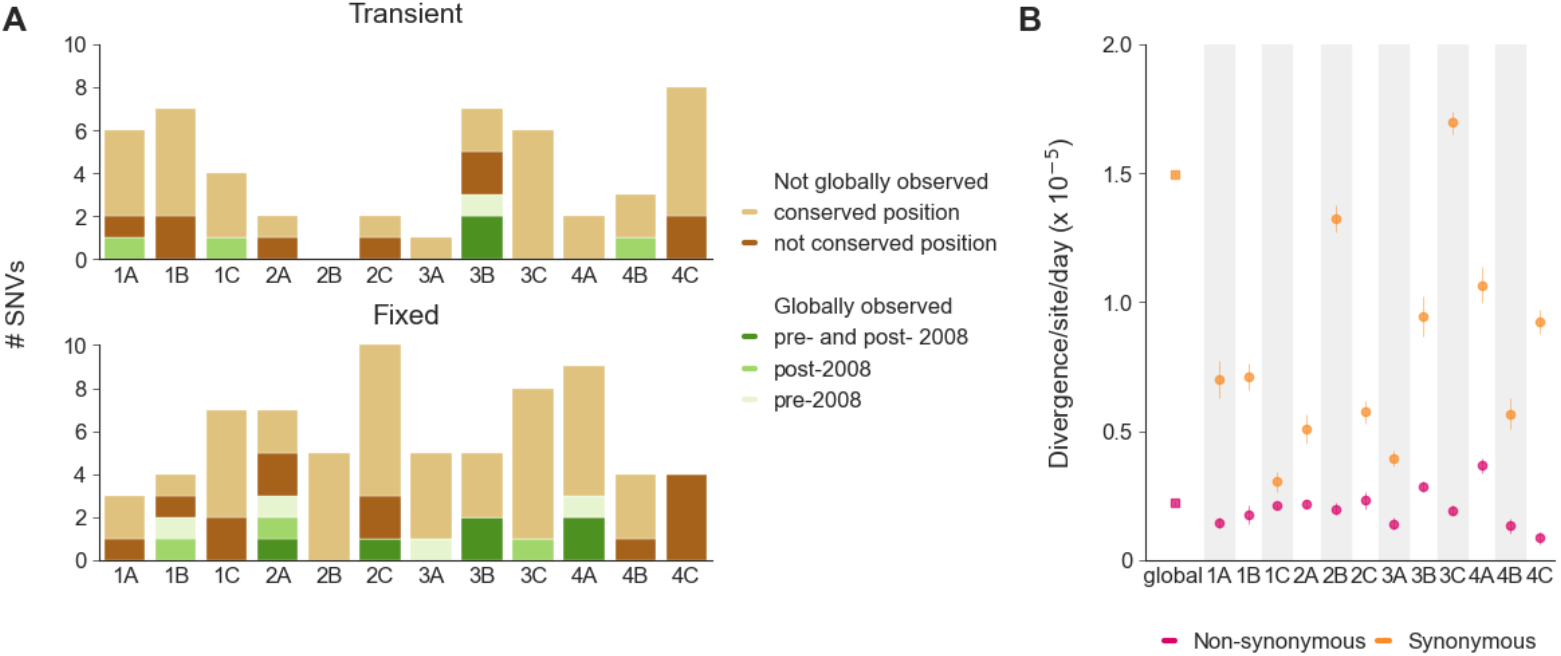
Experimental evolution vs global evolution. **A**, Number of non-drift-like AA mutations per population grouped by global observation patterns. **B**, Global and experimental rates of evolution at synonymous and non-synonymous sites within the PB2, PB1, PA, HA, NP, and NA genes. Experimental rates and global rates are shown as the mean and standard error calculated through linear regression.

As the starting virus was isolated in 2008 (see Methods), we also assessed if the observed non-drift-like AA mutations recapitulated global trajectories preceding and following virus circulation. Six populations acquired AA mutations that also arose in the global population after 2008 (Figure 6A; Extended Data Figure 7). Most notable was the HA mutation K92R (H3-numbering, antigenic site E) which fixed in population 2A and fixed in the global population in 2021. This mutation arose independently of mutation bias (SNV A323G resulted in the AA mutation) or linkage, in absence of immune selection, so its fixation cannot be attributed to mutation bias or linkage, and is therefore consistent with, though not demonstrative of, a fitness benefit independent of antigenic selection. Five populations showed reversion back to historical sequences, suggesting that the globally dominant AA is not fitness-optimal in this experimental system.

Lastly, we compared the evolutionary rates using experimental SNV rates versus global molecular clock rates for PB2, PB1, PA, HA, NP, and NA (Figure 6B). Non-synonymous rates were broadly similar overall, whereas synonymous rates were slower experimentally than globally across all populations except for population 3C. These slower rates may reflect weaker selection along with global epidemiological dynamics, as global synonymous changes likely often result from hitchhiking.

## Discussion

In this study, we assessed the predictability and repeatability of seasonal influenza A/H3N2 virus evolution by serially passaging the same A/H3N2 starting virus for ~1,200 generations in 12 independent lines from four HAE donors. We found that, each population evolved along unique trajectories and showed distinct dynamics, with minimal evidence for parallel evolution and no consistent pattern in which mutations arose within or across donors or segments, notwithstanding donor-level differences in the tempo of fixation and initial accumulation of nucleotide diversity. Compared to the evolutionary patterns of global A/H3N2 virus circulation in humans, our experimental populations generated AA mutations that were novel globally, occurred at globally conserved positions, predated global emergence, and reverted to historical sequences. These findings suggest that the evolutionary space of A/H3N2 viruses extends well beyond what is observed under natural conditions, and the observed natural trajectories only represent a small draw of possible evolutionary outcomes. In our system, virus evolution appears to be neither obviously repeatable nor meaningfully predictable.

Alternative evolutionary trajectories have been previously described for seasonal influenza A/H3N2 viruses, as the serially passaging of seasonal influenza viruses in both cell cultures as well as eggs have revealed HA changes that are distinct from those observed globally^32,33^. However, these changes are often linked to adaptations towards the different hosts environment. Conversely, in this study we employed a HAE system that recapitulates the cellular environment of the primary influenza infection site, and also found alternative trajectories when no obvious host adaptation was necessary. The stochastic diverging evolution observed here demonstrates that the accessible sequence space of human A/H3N2 viruses in absence of adaptive immune selection is potentially vast and under explored.

Beyond the existence of these alternative evolutionary trajectories, their character is also informative. In all populations we observed non-drift-like AA mutations at globally conserved positions, and for most genes these accounted for the majority of the observed mutations. A considerable proportion of these mutations were non-conservative (16/66, 8 fixed), and five populations reverted to AAs no longer observed in the global population. The non-neutral signal and the frequencies reached by these mutations is not consistent with negative selection acting on these positions, indicating that they are mutationally tolerant despite being near-invariant in nature.

This suggest that conservation of these positions may be maintained by the context in which the virus evolves, such as adaptive immunity, inter-host transmission bottlenecks, or competition between co-circulating lineages, rather than by intrinsic intolerance at the protein level.

Unique trajectories in sequence space are not a novel observation. Other long-term E&R studies have similarly reported that the majority of SNV and AA changes occurred within a single population^44^. However, despite following unique trajectories within sequence space, these studies found evidence of parallel evolution^39,42,43^, which we did not. One possible explanation is that, given the shorter time scale of our experiment, we simply have not observed this phenomenon yet. But this seems unlikely: the small genome of A/H3N2 viruses combined with its high mutation rate (~13.5 kb genome · ~3·10^−6^ mutations/site/generation), yields a substantial mutation supply. Assuming A/H3N2 viruses produce 10^8^ virion per 72-hours passage, this corresponds to roughly 100-fold the genome length in mutations per day. Signatures of parallel evolution would therefore have been expected to be detectable, making time an unlikely limiting factor.

However, a previous study investigating the long-term within-host evolution of seasonal influenza viruses during prolonged infections in immunocompromised individuals found evidence for parallel evolution, with mutations predominantly located in or adjacent to the antigenic sites of HA and the catalytic site of NA^45^. Although the human infections in that study represent the natural conditions under which influenza viruses replicate, the parallel antigenic evolution itself is evidence for directional selection enforced by the (compromised) adaptive immunity of the infected hosts. Given that our experimental systems lacked both an adaptive immune component and other obvious directional selective forces, the lack of parallelism is potentially unsurprising.

A more plausible explanation for the lack of parallelism might be the experimental set-up itself. The HAE system recapitulates the natural cellular environment to which our starting virus had already adapted over 40 years (1968 emergence to 2008 sampling). This contrasts with the other long-term E&R studies, which were conducted in laboratory environments unrepresentative of the organisms’ natural environment, and to which even laboratory-adapted strains had likely not fully adapted yet, particularly given the larger genomes and lower mutation rates of these organisms^46^. Indeed, several parallelism signatures observed in these studies have been shown to be beneficial under similar experimental conditions^42,43^. The lack of parallelism in our study may therefore reflect the fact that the experimental evolution closely models natural conditions to which the virus was already well adapted.

An important consideration when interpreting these results is the effective population size of our system. Although each passage generated a large virus population, the bottleneck of 1,000 infectious units imposed at every passage constrained the effective population size to the order of 10^3^. Selection was therefore only efficient for mutations with *s* greater than approximately 10^−3^, and both weakly beneficial and weakly deleterious mutations behave effectively neutrally. Two consequences follow. First, parallelism driven by weak-effect beneficial mutations would not be detectable here, and our parallelism results should therefore be read as applying to mutations with detectable non-neutral dynamics rather than to the full spectrum of beneficial mutations. Second, mildly deleterious mutations could drift to appreciable frequencies, which is why we restricted our interpretation of mutations at globally conserved positions to those showing non-neutral dynamics.

Although our experimental system modelled the natural environment of influenza A/H3N2 viruses in the absence of adaptive immune selective pressures, we cannot fully rule out the presence of selective constraints and experimental artifacts. Firstly, the HAE system maintained tissue-specific constraints, though they may vary by HAE donor. Secondly, the HAE cultures retained innate immune components (e.g., interferon signalling and mucociliary defences), which could still exert selective pressures. Additionally, we found evidence of bias towards increased AT content driven by elevated C:G→T:A and decreased A:T→G:C rates, consistent with previously reported C:G→T:A rates as the highest synonymous rates in global influenza virus data^47^ and similar to what is observed in the original LTEE^46^. Therefore, this bias may reflect influenza virus replication dynamics, though we cannot rule out that it represents an artifact of our experimental system.

This study has several additional limitations. We only assessed molecular evolution and thus lack the functional data and fitness data to link the genotypic evolution to the phenotypic evolution as well as assess the effect of (individual) mutations. Additionally, we had only four HAE donors, and although healthy, the donors were aged 41, 50, 63, and 73 years and 3 of 4 were male, making it impossible to assess the full range of donor-to-donor variability. Lastly, we assessed the evolution in the absence of antibody-mediated selection pressure and natural transmission fitness, both dominant forces in global influenza virus evolution. Therefore, we could not assess the repeatability or predictability in context of these constraints. Future experiments examining seasonal influenza evolution under adaptive immunity-mediated selection could offer insight in to repeatability and predictability of evolution reflective of natural conditions.

We showed that seasonal influenza A/H3N2 viruses evolving in a physiologically relevant cellular environment access a sequence space substantially larger than the one traversed by the global virus population. Parallelism between replicate lines was far from pervasive, suggesting that the virus’s potential evolutionary space is not intrinsically constrained to a small number of evolutionary paths. The trajectories we observed also included non-conservative mutations at positions held near-invariant globally and reversions to AAs the global population has long since abandoned, suggesting that the path taken by the global virus population may be constrained less by what the virus can tolerate than by the context in which it evolves. This combination of potentially large accessible sequence space and weaker than previously considered intrinsic constraint may contribute to the persistent challenge of predicting seasonal influenza evolution.

## Methods

### Selection of the virus strain

To ensure that our starting virus strain had a high fitness with a minimal number of deleterious mutations, we focused our selection on viruses positioned near the trunk of the phylogenetic tree for each gene segment. Here, we focused on the time window during the antigenic cluster transition from the Wisconsin/67/2005-like viruses to the Perth/16/2009-like and Victoria/208/2009-like viruses. These viruses circulated ~15 years before the study began and thus the natural evolutionary trajectory that followed has been documented. We selected A/Japan/WRAIR10159/2008 as our starting virus based on its whole genome sequence and proximity to the trunk of the phylogenetic tree from each segment. To create a minimally variable virus population, we produced our starting virus population using the recombinant methods described by Hoffmann et al^48^.

All experiments involving live, replication-competent influenza A/H3N2 virus were performed under Biosafety Level 3 (BSL-3) containment in accordance with institutional biosafety procedures and applicable regulations. All personnel involved in the laboratory work were vaccinated against seasonal influenza throughout the study. Appropriate procedures for the containment, handling, decontamination, and disposal of infectious material were followed throughout the experiments.

### Cell culture

MDCK cells were cultured in minimal essential medium (MEM no glutamine, Gibco), supplemented with 10% fetal calf serum (FCS; Sigma), 1% non-essential amino acids (NEAA; Gibco), 2 mM L-glutamine (Gibco), 100 units/mL penicillin, 100 units/mL streptomycin and 20 mM HEPES (Gibco) and incubated at 37°C and 5% CO_2_. Primary nasal human airway epithelial (HAE) cells were obtained from Epithelix (Switzerland) (Extended Data Table 3). Primary nasal HAE cells were maintained in PneumaCult-ExPlus medium (StemCell) in T75 flasks coated with type-I collagen (VitroCol, 5007-20ML, Advanced BioMatrix). Upon reaching 80–90 per cent confluency, cells were trypsinised and resuspended in PneumaCult-ExPlus medium to 0.4 µm pore-size transwell inserts (3470, Corning) which were coated with type-IV collagen (C7521-10MG, Sigma). When confluency was reached, apical media was removed, and basolateral media were replaced with PneumaCult-Ali medium (StemCell) to initiate differentiation at the air–liquid interface over 6 weeks. HAE cultures were assessed for differentiation by visual confirmation of beating cilia and mucus production. A subset of cultures was additionally cross-sectioned, H&E stained, and assessed by a histopathologist to confirm the presence of the expected differentiated cell types. Cells were washed once a week with HBSS for 10 minutes to remove mucus and cell debris. All cells were cultured at 37°C and 5% CO_2_ until experimentation.

### *In vitro* viral infection

For each passage, differentiated nasal HAE cells were inoculated with 1,000 infectious units of influenza A viruses by adding 100µL of 10^4^/mL TCID_50_ in HBSS to the apical side of the trans-well insert. Following inoculation cells were incubated at 33°C and 5% CO_2_ for 2 hours, after which the inoculum was removed and the apical side was washed twice with HBSS to remove unbound virus. After washing, the cells were returned to 33°C and 5% CO_2_ to incubate for a total of 72hrs. At 72 hours post-infection, the apical side of the culture was sampled by washing 2x100µL HBSS for 10 minutes to collect unbound virus. Viruses sampled after every 10^th^ passage were subject to sequence analysis.

### Viral titrations

Viral titrations were determined by infecting MDCK cells with ten-fold serial dilutions of virus stocks or HAE supernatants from every passage in minimal essential medium (MEM no glutamine, Gibco), 1% non-essential amino acids (NEAA; Gibco), 2 mM L-glutamine (Gibco), 100 units/mL penicillin, 100 units/mL streptomycin and 20 mM HEPES (Gibco) and 1 µg/mL TPCK-treated trypsin (Sigma–Aldrich). Cells were incubated at 37°C and 5% CO_2_ for 72 hours. After incubation, supernatant was removed and MDCKs were fixed with ice cold 80% acetone for 1 hour. Following fixation, plates were washed with TBS-0.05% Tween 20. Cells were incubated for 1 hour at room temperature with a monoclonal antibody (clone HB-65 IgG2a) directed against the nucleoprotein of influenza A virus (American Type Culture Collection, Manassas, VA) diluted in PBS containing 2% milk.

After incubation, plates were washed with TBS-0.05% Tween 20 followed by incubation for 1 hour at room temperature with goat anti-mouse IgG (H+L) Antibody, Human Serum Absorbed and Peroxidase-Labeled (SeraCare #5220-0341). After incubation, plates were washed with TBS-0.05% Tween 20. Development solution (0.1M NaAc and 0.1M citric acid) containing 1X 3,3′,5,5′-tetramethylbenzidine (TMB) solution and 1:3000 30% H_2_O_2_ was added, and the plates were developed at room temperature before adding stop solution (0.8M H_2_SO_4_). The optical density at 450 nm was measured using a Spectrostar nano BMG plate reader. Wells with an OD greater than 2-fold the mean OD of negative control cells were scored as positive. The frequency of positive and negative wells among the replicates per dilution was used to calculate the infectious virus titers (log TCID_50_/mL) according to the method of Reed and Muench^49^.

### RNA Extraction and polymerase chain reaction (PCR) amplification

Viral RNA was extracted from the HAE supernatant samples using the High Pure Viral RNA Kit (Roche, Basel, Switzerland) according to the manufacturer’s protocol and eluted in UltraPure DNase/RNase-Free Distilled Water (ThermoFisher Scientific, MA, USA). Viral RNA was amplified using one-step RT-PCR with primers targeted to the 3’ and 5’ conserved ends of the eight segments^50^. The primers include the CommonA-Uni12G (GCCGGAGCTCTGCAGATATCAGCGAAAGCAGG), CommonA-Uni12 (GCCAGAGCTCTGCAGATATCAGCAAAAGCAGG) and CommonA-Uni13G (GCCGGAGCTCTGCAGATATCAGTAGAAACAAGG)^50^. Briefly, RT-PCR reactions containing viral RNA, RT-PCR buffer, SuperScript III One-Step RT-PCR Platinum Taq HiFi DNA polymerase, RnaseOUT Recombinant Ribonuclease Inhibitor (Invitrogen, USA), dH_2_O and 20 µM of each primer were prepared^51^. Thermocycler conditions and ramp rates were based on the protocol from Van Poelvoorde et al.^51^. After amplicon generation, samples were purified and eluted in UltraPure DNase/RNase-Free distilled water using the MinElute PCR Purification Kit (Qiagen). The presence of the correct size PCR products was confirmed using a 1% agarose gel containing ethidium bromide (ThermoFisher Scientific) at 100V and visualised by the Vü Imaging System (Pop-Bio).

### Library preparation and next generation sequencing (NGS)

Library preparation and NGS were performed by GenomeScan B.V. (Leiden, the Netherlands). Sequencing libraries were prepared with NEBNext® Ultra II DNA Library Prep kit for Illumina (New England Biolabs) according to manufacturer’s instructions. Samples were then sequenced (1 Gb) on either the Illumina NovaSeq6000 or NovaSeq X platforms (as specified in Supplemental Table S1) to produce 2x150bp paired-end reads.

### Read processing

Raw reads were processed with a custom Snakemake-based ^52^ (v5.13.0) pipeline. In brief, reads were quality-trimmed using Trimmomatic^53^ (v0.39). Here, leading and trailing bases with a quality score (*Q*-score) below 3 were removed and an adaptive quality trimming (MAXINFO) was applied discarding reads < 30 bases and strictness of 0.4. Adapter sequences were removed using ILLUMINACLIP (TruSeq3-PE-2.fa:2:3:10). Reads were then aligned against the consensus coding domain sequence (cds) of the starting virus via Bowtie2^54^ (v2.4.5) using the very-sensitive-local settings. Only properly aligned paired reads with a mapping quality of at least 20 and a read length of at least 150 bases were retained. Coverage for each position in the cds was determined with a custom python script. Bases within each read with a *Q*-score below 30 were ignored. Final and intermediate read quality determined via FastQC^55^ (v0.11.9) as well as final genome wide coverage were assessed manually. Low quality (dips/gaps in segment coverage) and low coverage samples (median sample coverage <500×) were excluded from subsequent analysis and resequenced (as indicated in supplemental Table S1).

### Variant calling

Candidate SNVs were identified independently for each technical replicate population *p*_*r*_ and each timepoint *t* with a custom python script, adapted from Han et al.^56^. An SNV (denoted by *m*) was called if the frequency 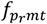 was ≥ 1%, the sequencing depth 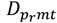 was ≥ 500× and the probability that the variant was the result of a base calling error (*p*_*Err*_) was < 0.1%^57^. The probability of a base calling error, *p*_*Err*_, was calculated using a binomial model, where the likelihood of observing at least 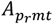 counts of the alternate allele of the total 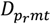 counts was modelled as^56,58^:

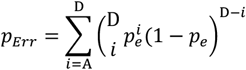

where 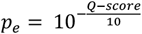.

SNV positions were annotated using a continuous numbering system that begins at the first nucleotide of the segment’s first ORF. Final SNVs were called if the SNV was observed in both technical replicates, SNVs only observed in one replicate were discarded.

We additionally evaluated whether an adapted version of the statistical framework described by Good, et al.^39^ (Supplemental Methods 1) could reliably discriminate true SNVs from sequencing errors. However, additional inclusion criteria were necessary to recover all SNVs observed above 10% (Supplemental Figure S3), given that under the employed quality criteria (500× coverage and *Q*30), the probability of such a variant resulting from error is *P*(*X* ≥ 50|*λ* = 500 · 00001) ≈ 10^−80^. Given this probability and the additional complexity introduced by those criteria, we concluded that the more parsimonious filtering approach described above was appropriate.

To estimate the true SNV frequency *f*_*pmt*_ we employed a maximum likelihood estimation (MLE) framework. Given the independent nature of technical replicates, the joint likelihood of observing 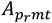 alternate allele counts at a total depth of 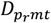 for replicates *r ∈* {1,2} follows a binomial distribution. We obtained the frequency estimate 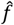 by minimising the negative log likelihood:

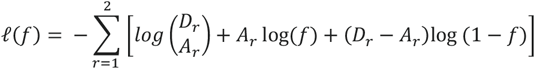

To account for sequencing noise and potential depth disparities across the replicates, optimization was performed numerically using the Nelder-Mead algorithm (SciPy^59^ *optimize* module (v1.9.3)) with the initial parameter estimate:

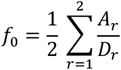

### Nucleotide diversity

Within-host genetic diversity was measured via nucleotide diversity ***π***. We calculated the nucleotide diversity for each population *p* at timepoint *t*, independently. For each site *l*, the per-site diversity is given by:

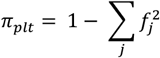

where *f*_*j*_ is the frequency of allele *j* at site *l* summed over all observed alleles at site *l* (including the ancestral allele)^60^. The whole-genome nucleotide diversity ***π*** was then calculated via:

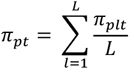

where *L* is the total cds length.

### Statistical comparison of nucleotide diversity over time between donors

To access differences in the initial rate of accumulation of whole genome nucleotide diversity between donors, we performed a linear mixed-effects regression on the nucleotide ***π***_*p*_ for each population *p* against the mean-centered timepoints *t*_c_, donor *d*, and their interaction term, with population included as a random effect. Here, we first identified the latest timepoint at which most populations still had a linear increase in nucleotide diversity by visually inspecting the nucleotide diversity per population plotted over time (Extended Data Figure 3). We included all timepoints up until this timepoint and centered on the mean of the included timepoints. We then determined the appropriate random-effects structure by comparing random intercept-only model with a random intercept and random slope model using a likelihood ratio test (LRT). The random intercept and slope model performed similar to the intercept-only model (*χ*^2^(2) = 1.51, *P* = 0.47), and we therefore continued with the intercept-only model. The models were implemented in R (v4.5.1)^61^ and fitted using the *lme4* package (v2.0.6)^62^. To test whether the rate of initial nucleotide diversity accumulation differed significantly between populations, we performed an *F*-test on the interaction coefficients. Finally, we performed pairwise comparisons on the estimated trends across donors using Tukey’s adjusting with the *emmeans* package (v2.0.4)^63^.

### SNV accumulation over time

The accumulation of SNVs over time for each population *p* was determined by calculating the total derived allele frequency as described by Good and colleagues^39^:

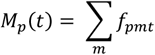

### Statistical comparison of SNV accumulation over time between populations

We estimated the rate of SNV accumulation over time per population *p* using a linear regression model on total derived allele frequencies *M*_*p*_(*t*) against the mean-centered timepoints *t*_c_, population *p* and their interaction. The model was implemented in R (v4.5.1)^61^. To test whether the rate of SNV accumulation differed significantly between populations, we performed an *F*-test on the interaction coefficients. The comparison was performed on the total derived allele frequencies calculated from all SNVs, non-synonymous SNVs only, and synonymous SNVs only.

### Statistical comparison of SNV accumulation over time between donors

To infer if there were significant differences in the rate of SNV accumulation between donors, we performed a linear mixed-effects regression on total derived allele frequencies *M*_*p*_(*t*) against the mean-centered timepoints *t*_c_, donor *d*, and their interaction term, with population included as random effect. We first determined the random-effects structure by comparing a random intercept-only model with a random intercept and slope model using an LRT. The random intercept and slope model had a significant better fit for all SNVs (*χ*^2^(2) = 152.76, Δ Akaike information criterion (AIC) = 148.76), non-synonymous SNVs only (*χ*^2^(2) = 86.86, Δ AIC = 82.86), and synonymous SNVs only (*χ*^2^(2) = 176.3, Δ AIC = 172.3) (all *P* < 2.2 × 10^−16^). However, the random intercept and slope model resulted in convergence issues (singular fit). We therefore simplified the random effects by setting the correlation between the random intercepts and slopes to zero. Because uncorrelated random-effects are not invariant to shifts in the predictor^62^, we changed the mean-centered timepoints *t*_*c*_ to the raw timepoints *t*. Model performance was evaluated again using an LRT. The uncorrelated version of the random intercept and slope model still provided a significantly better fit compared to the intercept only model (all SNVs: *χ*^2^(2) =143.88, Δ AIC (relative to intercept only model) =141.88; non-synonymous SNVs only: *χ*^2^(2) =83.51, Δ AIC =81.51; and synonymous SNVs only: *χ*^2^(2) =168.5, Δ AIC =166.5; all *P* < 2.2 × 10^−16^). For the non-synonymous SNVs only model the convergence issues could not be resolved using the uncorrelated version. Yet we choose to use this model as it was the more appropriate design for our experimental set-up. The models were implemented in R (v4.5.1)^61^ and fitted using the *lme4* package (v2.0.6)^62^. We tested if the rates of SNV accumulation differed between donors via Kenward-Roger^64^ *F*-tests on the interaction term. We have reported the results of the uncorrelated random effects in the *Results* section above; all conclusions hold using the model with correlated random effects.

### Phylogenetic inference

Maximum likelihood (ML) phylogenetic trees were reconstructed independently from the consensus cds sequences using IQ-TREE 2^65^ (v2.2.0.3) with the GTR+G4 nucleotide substitution model and rooting on the starting virus consensus cds.

### Within-host simulations

To test whether our experimental system contained only neutral evolutionary forces, i.e. genetic drift, or whether there was evidence of non-neutral forces, we devised an *in silico* model that closely mimics our experimental set-up but where we ensured mutation survival was only the result of genetic drift. This model comprises three components: an infection, a growth, and a sampling component.

The infection component mimics the infection of the HAE cultures. Infection was initiated with *V*_*input*_ infectious units of which we assume a fraction *f*_*si*_ will be successful infecting the (HAE) target cells *C*, resulting in *V*_*infect*_ infecting virions. We set *V*_*input*_ to 1000 virions in accordance with the number infecting units used in the experiment.

The growth component mimics the viral growth within the HAE culture during the 72 hours incubation period. Here we computed the changes in virus population size *V* and available target cells *C* using a target-cells limited model^56,66^. During the incubation period, new virions are produced at a rate of *βCV*, where *β* denotes the per-cell per-virion contact rate. Each infected cell will produce *r* virions before it is rendered unproductive. We computed the viral growth during the incubation period by numerically solving the following systems of ODEs:

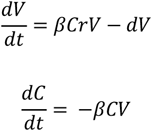

where *t* is a replicate generation. We computed *β* by fixing the within-host basic reproduction number *R*_0_ and the initial target cell population size *C*_0_:

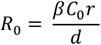

For every generation *t*, we tracked the frequency distribution of genotypes present. We assumed that the experiment starts with a binary genome of length L, where each locus is neutral throughout. At each generation, the number of virions acquiring a random single locus mutation follows a Poisson distribution with mean *V*(*t*)*µL*. The genotype distribution for the next generation *t* + 1 was then obtained by Poisson resampling individuals with mean *V*(*t* + 1)*f*_*g*_ (*t*), where 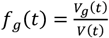 is the frequency of genotype *g* at generation *t*.

Finally, the sampling component mimics how we sampled for sequencing and inoculation of the next passage. After the growth, we assumed that only a fraction of the virion population *f*_*apical*_ is released into the apical compartment of the transwell plate. We obtained the genotype distribution of the virions released apically *V*_*apical*_ = *V* · *f*_*apical*_ by Poisson resampling of the total virion population according to the genotype frequencies *f*_*g*_. Next, we modelled the number of virions that would be recovered from performing apical washes by assuming that a fraction *f*_*wash*_ of the apical virion population is recovered, giving *V*_*wash*_. We obtained the genotype distribution of these virions by Poisson resampling *V*_*apical*_ according to the genotype frequencies *f*_*g,apical*_. From *V*_*wash*_, we sampled 1/4 of the sample for sequencing, giving *V*_*seq*_ = 0025 · *V*_*wash*_, with genotype distribution obtained via Poisson resampling according to *f*_*g,wash*_. This sequencing sample was then split into two equal technical replicates. The first replicate 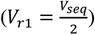 was assigned a genotype distribution by Poisson resampling of *f*_*g,seq*_. The genotype counts of the first replicate were then subtracted from the sequencing counts, *N*_*g,r*2_ = *N*_*g,seq*_ – *N*_*g,r*1_, from which the genotype frequencies for the second replicate, *f*_*g,r*2_ were obtained by normalising *N*_*g,r*2_.

From the remaining wash sample, we took 1/3 of the sample for inoculation of the next passage, giving 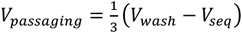. The genotype counts of the remaining wash population were calculated by subtracting the sequencing counts from the wash counts, *N*_*g,remaining*_ = *N*_*g,wash*_ – *N*_*g,seq*_. The corresponding genotype frequencies *f*_*g,remaining*_ were obtained by normalising these counts. The genotype distribution of *V*_*passaging*_ was then obtained by Poisson resampling according to *f*_*g,remaining*_ This sample was subsequently resampled to *V*_*input*_ virions, with the genotype distribution of the inoculum obtained by Poisson resampling according to *f*_*g,passaging*_.

To mirror our experiment, we simulated 100 passages. For each passage we performed the same sampling procedure. However, to obtain frequency trajectories with the same timing as the actual experiment, we only ‘sequenced’ both technical replicates of every tenth passage. To obtain at least 20000 simulated SNV trajectories, we ran 100 simulations per variable L for the cds lengths of the PB2, PB1, PA, HA, NP, and NA segments, and 200 simulations for the M and NS segments. We tested variable sets for *f*_*apical*_ and *f*_*wash*_, which showed robust results (Supplemental Figure S4). All parameter values used in the final model are given in Supplemental Table S2.

### Hidden Markov model to estimate the approximate appearance time and fixation of SNVs

To track the emergence, fixation, and loss of SNVs within each population we utilised a hidden Markov model (HMM)^39^ that aggregates information across timepoints. In this HMM, all SNVs start in the ancestral state ***A*** where *f*_*pmt*_ = 0. At each timepoint there is a finite probability that the mutation transitions from the ancestral state ***A*** into a polymorphic state ***P*** or a fixed state **F**. In our model we allowed SNVs to directly transition from ***A*** to ***F***, as the large interval between windows allowed for unobserved dynamics during which an SNV could have emerged and become fixed without passing through an observed polymorphic state. From the polymorphic state ***P***, SNVs can transition back to ***A*** (go extinct) or to **F** or remain in ***P***. From the fixed state ***F***, SNVs can transition back to ***P*** or directly transition back to state ***A***, once again accounting for potentially unobserved dynamics. The schematic of the Markov chain is illustrated in Supplemental Figure S5.

We estimated the initial transition probabilities empirically, by random subsampling 30% of the observed SNV trajectories, assigned each timepoints to ancestral ***A*** (*f*_*pmt*_ ≤ 0001), fixed ***F*** (*f*_*pmt*_ ≥ 0099), and polymorphic ***P*** states via frequency threshold, and applying Laplace smoothing (*α* = 1) to all permissible transitions to account for unobserved transitions. This gave us the following transition rates:

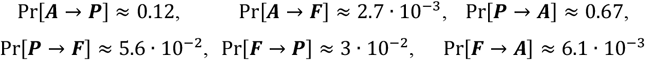

though the results are robust to the initial transition probabilities.

The HMM accounts for sequencing noise by modelling the alternate allele counts *A*_*pmt*_ as binomially distributed^39^, evaluating continuous emission likelihoods via regularised incomplete beta functions. We calculated log-emissions for the ***A*** and ***F*** states as well as error vectors 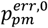 and 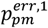, using the total allele counts *D*_*pmt*_, the alternate allele counts *A*_*pmt*_ and the ancestral allele counts *R*_*pmt*_. Here, 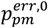 represents the SNV-specific false-positive error rate of observing alternate reads within the ***A*** state, while 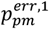 is the false-negative error rate of observing ancestral reads within the ***F*** state. These error profiles were updated iteratively at each maximization step by weighing *D*_*pmt*_ by posterior state densities derived via the forward-backward algorithm. After this process was repeated for 1,000 iterations (sufficient for convergence), we applied a standard Viterbi pass to obtain the most-likely state trajectory *L*_*pmt*_.

Given that we had two technical replicates and that these two replicates reflected the same underlying state, we pooled the data across replicates by summing their respective counts. Specifically, we used the total alternate allele counts 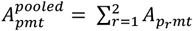 and the total allele counts 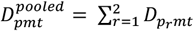 as the combined emission observations per SNV.

### Statistical comparison of first fixation event between donors

For each population *p* we identified the first timepoint *t* at which any SNV *m* had fixed, *TF*_*p*_. Let *S*_*p*_ denote the set of all SNVs detected in population *p*. We then obtained the set of SNVs that have fixed in population *p* via:

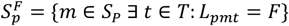

where *T* is the set of all timepoints. For each SNV in the set 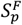 we obtained the earliest fixation event via:

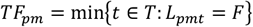

We then obtained *TF*_*p*_via:

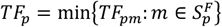

We evaluated difference in first fixation timing between donors using a gamma generalised linear model (GLM) with a log link. The model was implemented in R (v4.5.1)^61^. Overall donor effects were evaluated via an Analysis of Deviance (F-test) comparing the model to an intercept-only null, using Pearson’s chi-squared (*χ*^2^) dispersion. We performed a post-hoc exploratory contrast to compare the mean timing for the first fixation event of donors 3 and 4 with that of donors 1 and 2. The contrast’s *P*-value and 95%-CI were Scheffé-adjusted. Finally, pairwise donor comparisons were performed using Wald’s t-tests with Holm correction.

### Statistical identification of non-drift-like SNVs

To identify SNV trajectories that deviated from neutral expectation (drift), we devised a statistic framework that utilised in the max SNV frequency as well as the most-likely sequence of states *L*_*pmt*_ to assess if the SNV significantly differed from our drift-like null-model trajectories. The simulated trajectories from our within-host model serve as the null model for this comparison.

We quantified whether an observed SNV trajectory deviates from the null model using a composite test statistic that summarises evidence across two metrics that we expect to differ between drift-like (neutral) trajectories and non-drift-like trajectories: the max frequency, and transit time^39^. For both these metrics we extract information from the most-likely states *L*_*pmt*_. Over time, each SNV can have one or multiple observations, where the number of observations is defined as:

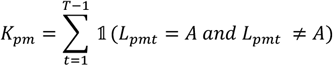

where *T* is the set of all timepoints. The start of these observations is given by:

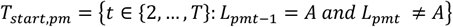

For each observation *k*, the end point of the observation is defined as:

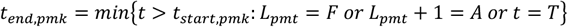

where the *k*-th observation corresponds to the *k*-th element of *T*_*start,pm*_. The point at which the achieved its highest frequency during that observation is simply given by 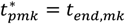 if the SNV fixed during the observation or by:

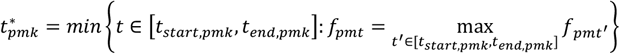

if the SNV did not reach fixation during that observation. The max frequency of that observation *k* is given by:

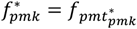

For non-drift-like trajectories we expect higher frequency than those reached as a result of chance. The transit time is the time from emergence to max frequency and can be obtained via:

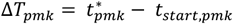

For non-drift-like trajectories we expected a shorter transition time as neutral trajectories will only gradually increase in frequency.

For each observation *k* for each SNV *m* in each population *p*, we sampled 10000 observations from the simulated trajectories (L = cds of the segment of the observed SNV), and numerically calculated a single-statistic P-value comparing the max frequency 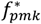 and the transit time Δ*T*_*pmk*_ :

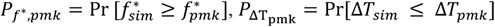

We then combined these single-statistic P-values into a composite statistic^39^:

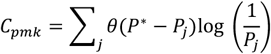

where *P*_*j*_ indicates the single-statistic *P-*values in 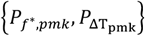, *θ*(·) is a Heaviside step function, and 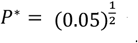, which ensures that the statistic is dominated by largest deviations in one of the statistics, rather than smaller or moderate deviations in all of them. For this composite statistic we calculated the composite p-value by sampling another 10000 observations from the simulated trajectories:

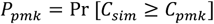

After estimating a *P*-value for every observation, we corrected for multiple hypothesis testing across all populations, SNVs and observations, via the Benjamini-Hochberg (BH) procedure^69^, for which we selected a false discovery rate (FDR) of 0.01 based on a comparison across candidate values (Supplemental Figure S6), to include the most true set of non-drift-like signatures.

Finally, we assigned each SNV one of three evolutionary status classifications: drift, transient, and fixed. Drift SNVs are SNVs that were not significantly different from the simulated trajectories, whereas both transient and fixed SNV were significantly different (i.e. non-drift-like). Here, we chose to separate the non-drift-like SNVs into separate classifications, transient and fixed, as we did not observe fixation in any of the simulated trajectories (Figure 3A) and therefore fixation itself is evidence against drift.

### Haplotype reconstruction

We reconstructed the most parsimonious viral haplotype for each segment from all non-drift-like SNVs in each sample using the ML approach developed by Ghafari et al.^70^ and implemented as described by Han, et al.^56^. Briefly, a list of candidate full haplotypes was generated from all combinations of observed SNVs with a frequency ≥5% (non-drift and fixed SNVs only, unless specified differently) across the segment. To ensure that haplotypes supported by physical linkage (co-mutation) on single reads were prioritised over those containing rare, independent SNVs, each potential haplotype *h* was scored according to:

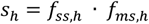

where *f*_*ss*.*h*_ and *f*_*ms*.*h*_ are both joint probabilities for the haplotype *h. f*_*ss,h*_ is the expected joint probability frequency calculated from the observed independent SNV frequencies found in haplotype *h* and *f*_*ms*.*h*_ is the multi-site joint probability derived from the observed frequencies of reads spanning the highest hierarchical combinations of co-mutational sites.

Assuming the true underlying viral population consists of set of *H* haplotypes with frequency *q*_*h*_, the expected partial haplotype frequencies *q*_*k*_ for a given read type *k*, defined as the unique tuple of polymorphic sites covered by a read, can be modelled as:

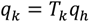

where *T*_*k*_ is a projection matrix mapping full haplotype frequency onto the specific sub-allele observed in reads spanning one or more polymorphic sites.

Beginning with *H* = 1, the optimal full haplotype frequency *q*_?_ is inferred by maximising the likelihood function:

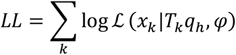

where *ℒ* is Dirichlet multinomial likelihood, *x*_*k*_ is the observed variant count data for read type *k*, and *φ* overdispersion parameter.

We estimated a global value for *φ* from the variance observed between the two independent technical replicates for each SNV in total set *M* of all identified SNVs using an MLE framework. For each SNV *m* in population *p* at timepoint *t*, we modelled the joint likelihood of observing 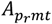 alternate allele counts at a total depth 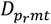 for replicates *r ∈* {1,2} using a beta-binomial distribution with the frequency estimate 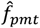 as the mean and concentration parameter *θ* = (1 ™ *φ*)/*φ*, to account for additional binomial variance beyond that expected under the binomial model. We then obtained *φ* by minimising the negative log likelihood:

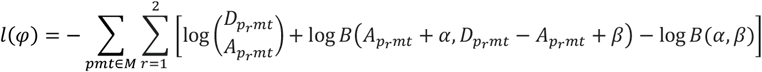

where 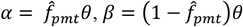, and *B*(·) represents the beta function. From our replicate data, we estimated *φ* ≈ 1 · 10^−3^.

Haplotype frequency optimization was performed via simulated annealing by running two independent searches for a minimum of 5000 iterations until convergence. At each iteration, the candidate frequency distribution for *q*_*h*_ was sampled from a Gaussian distribution centered at the distribution of the previous state with a standard deviation of 0.05. One additional haplotype from the candidate list was introduced during each successive round of optimization (*H* + 1). Any SNVs in the observed partial haplotypes that were unaccounted for in the current set of *H* full haplotypes were assumed to be generated from a cloud of “noise” haplotypes that were present in no more than 1%. The final most parsimonious number of haplotypes (*H*) was determined by selecting the model that minimises the Bayesian Information Criterion (BIC). Reconstruction was performed independently for each technical replicate; final haplotype frequencies were then estimated as the mean across the two.

### Statistical identification of potential hitchhiking and cooccurring SNVs

For the non-drift-like SNVs we determined whether the SNV’s trajectory could be explained by its putative linkage to another SNV. For this, we defined two categories of linked SNVs pairs. Hitchhikers consist of a non-synonymous (the putative driver) and synonymous (the putative passenger) SNV, where the passenger’s trajectory is hypothesised to reflect its linkage to the driver with a putative effect on fitness. Cooccurring pairs consist of either two non-synonymous SNV or two synonymous SNVs, where the trajectory of one may be driven by the other, or where there is a putative combined epistatic effect on fitness.

Putative linkage pairs were identified by computing the Pearson correlation between the trajectory of frequency changes (Δ*f*_*pm*_) for each SNV. To determine whether the observed correlation exceeded what would be expected by chance, we permuted the frequency trajectory of the driver (or first SNV in the cooccurring pairs) for 10,000 iterations, computing the Pearson correlation coefficient between the permuted trajectory and unshuffled passenger (or second SNV) trajectory at each iteration. The resulting distribution of *r* values served as the null model from which we calculated an empirical *P-*value:

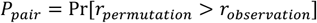

A pair was retained if the Pearson correlation between the Δ*f*_*pm*_ trajectories exceeded *r*_*observation*_ > 009 and *P*_*pair*_ < 0005. For pairs with SNVs located on the same segment we reconstructed the haplotypes over time (as described above) tracking the two SNVs, we then calculated the normalised linkage disequilibrium (*D*^’^; described below) to determine actual genetic linkage. We set *D*^’^ threshold ≥0.95 and labelled the confidence level for pairs that met this criterion as strong. The confidence level for pairs that did not meet this criterion or with SNVs on different segments was set to putative.

### Linkage disequilibrium

We computed the normalised linkage disequilibrium for hitchhikers and cooccurring pairs located on the same genome segment following Lewontin^71^. For each pair, we focused on the earliest timepoint at which both SNVs were simultaneously observed in a polymorphic state (i.e. *L*_*pmt*_ = *P* both SNVs). We first computed the linkage disequilibrium (D) via:

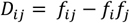

where *f*_*i*_ and *f*_*j*_ are the site-independent frequencies for sites *i* and *j*, respectively, and *f*_*ij*_ is the estimated haplotype frequency. The normalised linkage disequilibrium (*D*^’^) was then calculated by dividing *D* by its theoretical maximum *D*_*max*_:

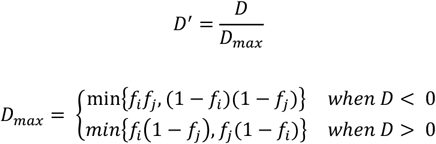

### Inference of mutation bias

To assess mutation bias within each experimental population, we first calculated the transition transversion (Ti/Tv) ratio from all synonymous SNVs within the non-overlapping ORFs (PB2, PB1, PA, HA, NP, NA). We then assessed whether this estimated ratio differed significantly from the expected Ti/Tv ratio of ~3^40,41^ via two-tailed binomial tests^46^ per population. As we observed a similar trend across populations, we used a binomial generalised linear mixed-effect regression model (GLMM) with a random intercept per population to estimate the experiment-wide Ti/Tv ratio. The estimated ratio and corresponding 95%-CI were obtained by exponentiating the log profile likelihood CI. We then repeated this analysis for the entire genome by including the synonymous SNVs located within the M1/M2 and NS1/NEP overlapping ORFs, to confirm that the results hold. For this analysis we still excluded the alternative PB1-F2 and PA-X ORFs given their low and strain-dependent expression levels^72^. To confirm whether the estimated Ti/Tv ratios were not an attribute of ancestral sequence composition we also calculated the expected Ti/Tv under neutral expectation (i.e. 0.5 as there are twice as many transversion pairs as there are transition pairs).

To identify which specific substitution types drove the observed change in Ti/Tv ratio, we decomposed the (synonymous) SNVs observed across the non-overlapping ORFs into six canonical substitution types (A:T→T:A, A:T→C:G, A:T→G:C, C:G→T:A, C:G→A:T, and C:G→G:C). To test whether the mutational spectra varied significantly across populations we performed a Monte Carlo-based Pearson’s chi-squared test using 10,000 simulated replicates. As mutational spectra did not vary significantly, we pooled the observed counts. We then calculated the expected mutational spectrum based on the ancestral opportunity weighted according to the expected Ti/Tv ratio of 3^73^ and the estimated Ti/Tv ratio of 11.75 and scaled by total number of synonymous SNVs observed. Using two-tailed binomial tests^46^, we then evaluated whether observed counts deviated significantly for the expected, with *P*-values adjusted for multiple hypothesis testing using the Holm method. We repeated this analysis for the individual populations and non-overlapping ORFs, and found that that increased bias towards C:G→T:A mutations holds. The GLMM and chi-squared test were implemented in R (v4.5.1)^61^, using the *lme4* package (v2.0.6)^62^ to fit the GLMM.

### Inference of codon usage bias (CUB)

We inferred if there was evidence for CUB within each population by assessing the directional associations between synonymous SNVs and its effect on codon usage relative to the preferred codons in the ancestral sequence. To do so, we first established the preferred codon for each AA in the ancestral sequence, i.e. the codon with the highest frequency. Based on this baseline SNVs were assigned to one of three categories: “toward preferred” if the mutated codon matched the preferred codon; “from preferred” if ancestral codon matched the preferred codon but the mutated codon did not; or “neutral” if neither the ancestral or the mutated codon matched the preferred codon. The proportional distribution of these categories was then quantified for each population across all synonymous SNVs, as well as stratified by classification (i.e. drift, transient, and fixed).

### Identification of parallel evolution signatures

To assess whether there was evidence for parallel evolution within our experimental system, we implemented a permutation-based parallelism test evaluating whether mutations occurred at the same genomic positions across populations more often than expected by chance. For this analysis we initially focused on non-drift-like non-synonymous SNVs located within the PB2, PB1, PA, HA, NP, and NA viral genes, but later also ran separate analysis for the overlapping ORFS pairs of M1/M2 and NS1/NEP as well as non-drift-like synonymous SNVs.

For each viral gene, the number of available sites was estimated per substitution type as described above. We then counted the total and unique non-drift-like SNVs per gene and substitution type, as well as the number of populations in which each unique SNV occurred (multiplicity). We then simulated 10,000 draws by randomly redistributing each population’s observed SNV counts across the available sites. We then quantified parallelism via two complementary statistics: a site-count statistic, defined as a site with a multiplicity of two or more; and a multiplicity-weighted statistic, defined as the sum of per-site multiplicities. For both statistics we then calculated an empirical *P*-value:

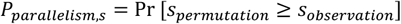

where *s* is the statistic evaluated.

### Comparison with global influenza A/H3N2 virus surveillance data

To compare how our evolutionary trajectories deviated from the global evolutionary trajectories, we first downloaded all available human lineage influenza A/H3N2 virus sequences collected from human hosts between 1968 and December 31^st^ 2025 and submitted before May 13^th^ 2026 from the Global Initiative on Sharing All Influenza Data (GISAID) EpiFlu database^74^. For each genome segment, we first removed low quality (>1% ambiguous nucleotides) sequences, we then translated the cds to the ORFs of the genes encoded on the segment. For each AA position within each gene, we then calculated the AA frequencies over time, binning sequences into time windows of variable sizes depending on data availability. For the earliest decades (1968-1997), sequences were grouped into five-year windows to ensure sufficient sampling depth. From 1998 onwards, each calendar year was treated as an independent window. To minimise the contribution of substitutions arising from passage adaptation, we also applied passage history filters based on data availability. We included all available data from windows between 1968 and 1999. For the 2000-2009 windows, we included clinical and cell-passaged sequences, and from 2010 onwards only clinical sequences. The exact number of sequences included per time window per segment are provided in Supplemental Table S11.

For each non-drift-like AA mutation, we determined whether it occurred at a global conserved position, defined as a site where a single AA maintained a frequency ≥ 90% in all time windows, and whether it was observed globally both pre-and post-2008, defined as mutated AA reaching frequency ≥ 10% in a time window.

### Within-host evolution rates

For each population *p* and each substitution type *s*, we estimated the within-host evolution rate as the slope of a linear regression of *M*_*ps*_ (*t*)/*n*_7_ against time in days. Here, *M*_*ps*_ (*t*)/*n*_*s*_ is the total derived allele frequency, i.e. the sum of SNVs frequencies, for all SNVs of type *s* in the PB2, PB1, PA, HA, NP and NA ORFs for population p at timepoint *t*, and *n*_*s*_ is the number available sites for type *s* within these ORFs^73^. Time in days is obtained for each timepoint by multiplying the passage number by three (72 hours per passage).

### Global rates of evolution

To estimate the global rate of evolution we used an adapted version of the molecular clock approach as described by Xue and Bloom^73^. To do so, we first randomly down-sampled our downloaded GISAID sequences to 100 sequences per year, prioritising full cds length and clinical sequences where data permitted. We focused on the time window post the collection date of our starting virus (A/Japan/WRAIR10159/2008; 2008-05-11).

Synonymous and non-synonymous SNVs relative to the starting virus were then identified and summed across the PB2, PB1, PA, HA, NP, and NA genes. Finally, a linear regression model of cumulative divergence per site per days elapsed relative to the starting virus’s collection date was fitted separately for each mutation type to determine the global rate of evolution.

## Acknowledgements

We would like to thank all authors and contributors of the GISAID EpiFlu database for sharing their influenza A/H3N2 sequence data.

## Author contributions

CAR and AXH contributed to conceptualization. AJHR, KDH, ZFG, DE, AXH, and CAR contributed to methodology. KDH, SL, KH, LEG, MH, and ZFG executed the experimental work. AJHR and AXH contributed to software, formal analysis, and data curation. AJHR, KDH, AXH, and CAR contributed to validation and visualization. CAR was responsible for funding acquisition. AJHR, KDH, and CAR wrote the original draft. All authors contributed to review & editing. All authors read and approved the final version of the manuscript.

## Competing interests

We declare no competing interests.

## Data and code availability

All raw sequence data are accessible at the NCBI sequence read archive under BioProject ID PRJNA1549747, available via: https://www.ncbi.nlm.nih.gov/bioproject/1549747. All analysis scripts and notebooks are at available at https://github.com/AMC-LAEB/LTEE-flu. All computational analyses were performed in python (v3.10.6) unless stated otherwise. Processed data required to replicate the results have been archived on Zenodo and are available via: https://doi.org/10.5281/zenodo.23661876.

## Funding

This work was supported by the European Research Council (ERC) Consolidator Award under grant number 818253.

## Extended data

**Extended Data Figure 1.**
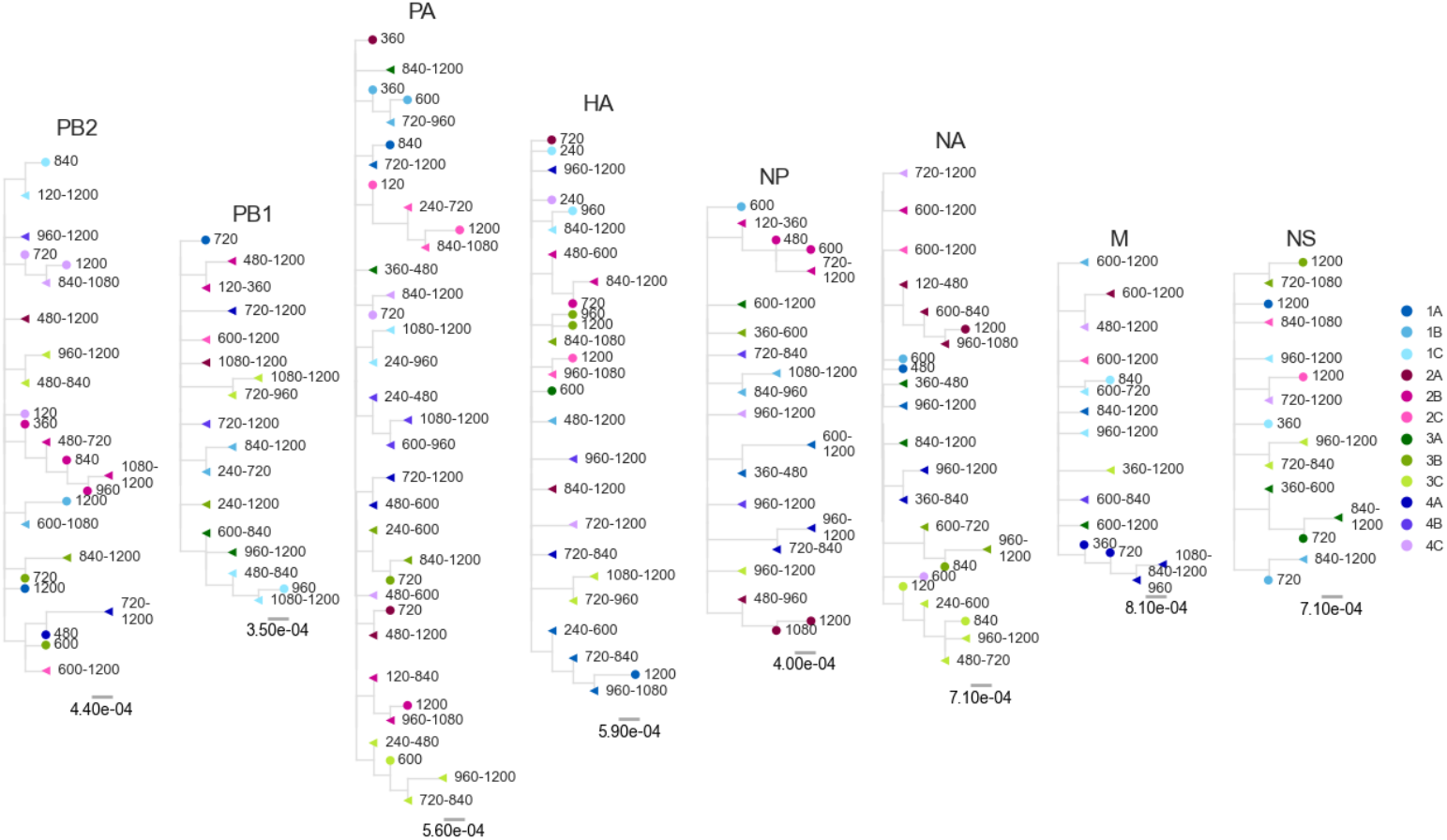
Segment specific phylogenetic trees from the population consensus sequences at each sampling timepoint (in generations).

**Extended Data Figure 2.**
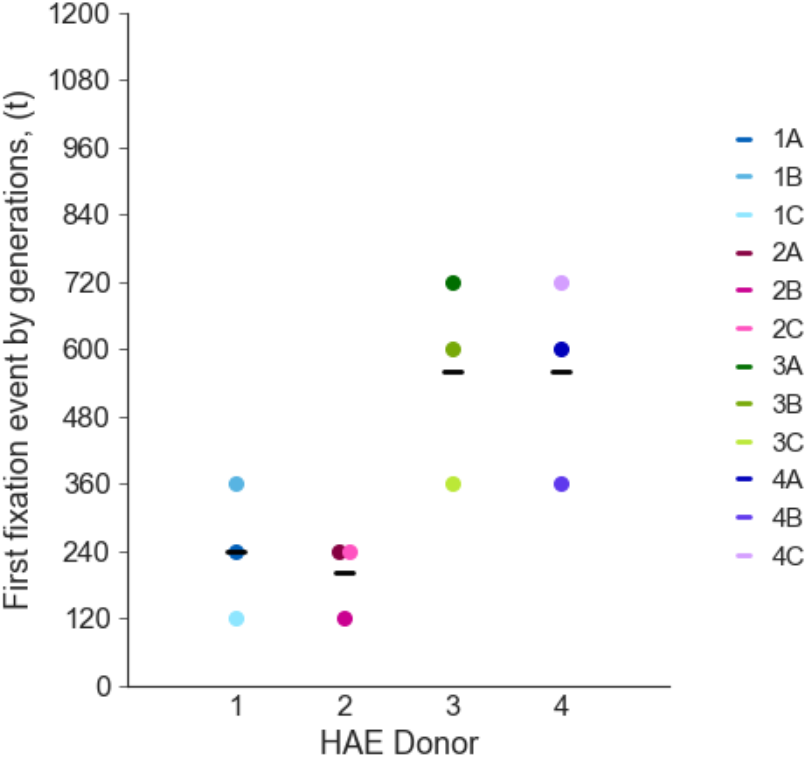
Timing to first fixation event per HAE donor. Coloured points represent the independent experimental population. Horizonal black bars indicated the donor mean timing to first fixation event per donor.

**Extended Data Figure 3.**
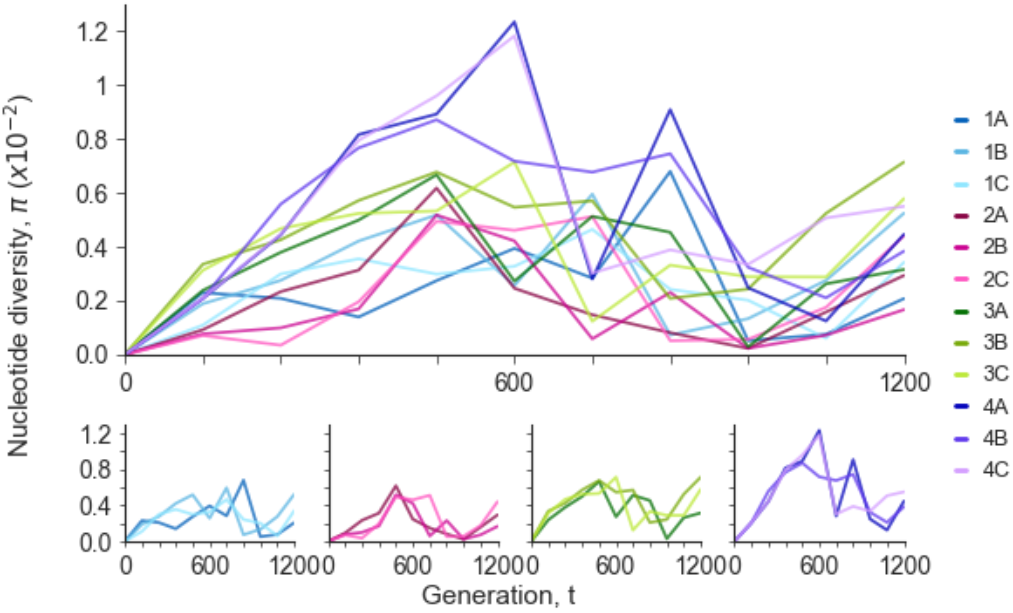
Genetic diversity over time measured by nucleotide diversity *π*. Top panel shows all populations, whereas bottom panels are separated by donor.

**Extended Data Figure 4.**
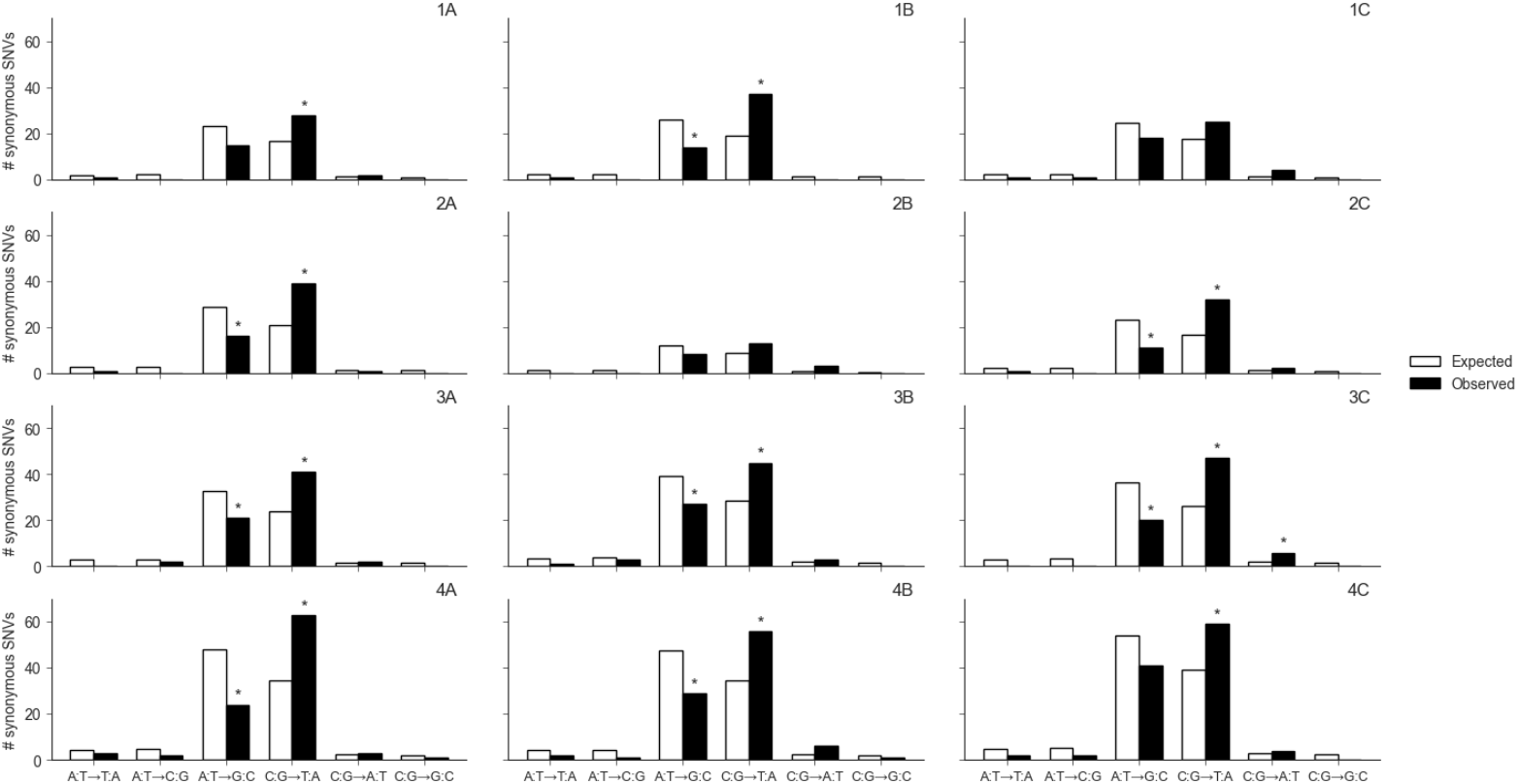
Expected and observed mutational spectra for synonymous SNVs per population. All synonymous SNVs in the non-overlapping ORFS of PB2, PB1, PA, HA, NP, and NA were included. Expected spectrum was calculated from the ancestral opportunity scaled by the total number of synonymous SNVs in the population, weighted according to the expected Ti/Tv ratio of 3. Asterisk indicates Holm-adjusted *P*<0.05(two-side binomial tests). Conclusions hold under the mean population-wide estimated Ti/Tv ratio of 11.75.

**Extended Data Figure 5.**
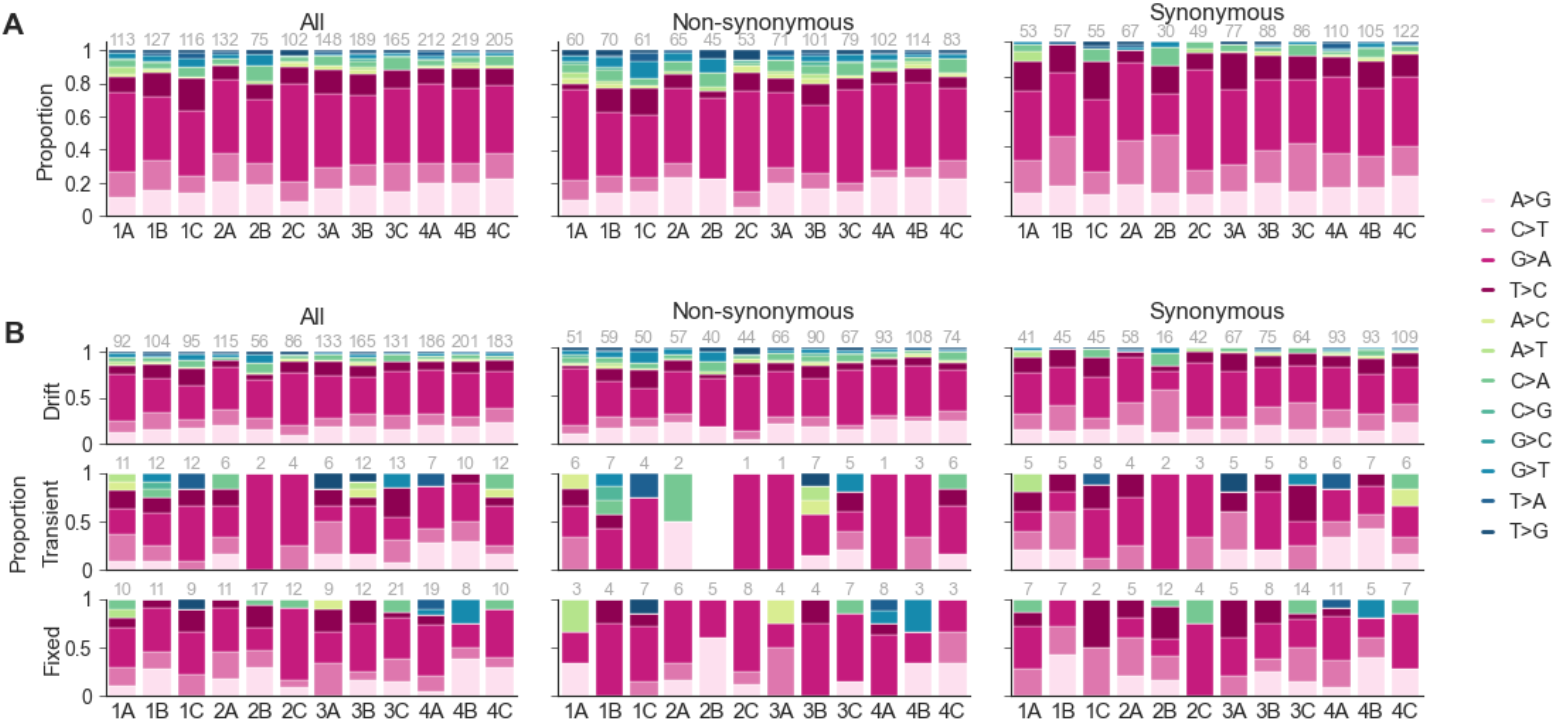
Proportion of nucleotide changes per population. **A**. Proportions of changes for all, non-synonymous, and synonymous SNV per population. **B**. Proportion of changes for all, non-synonymous, and synonymous substitutions separate by SNV classification. Pink shades indicated transitions and the blue green shades indicated transversions. The number above each bar indicates the number of SNVs.

**Extended Data Figure 6.**
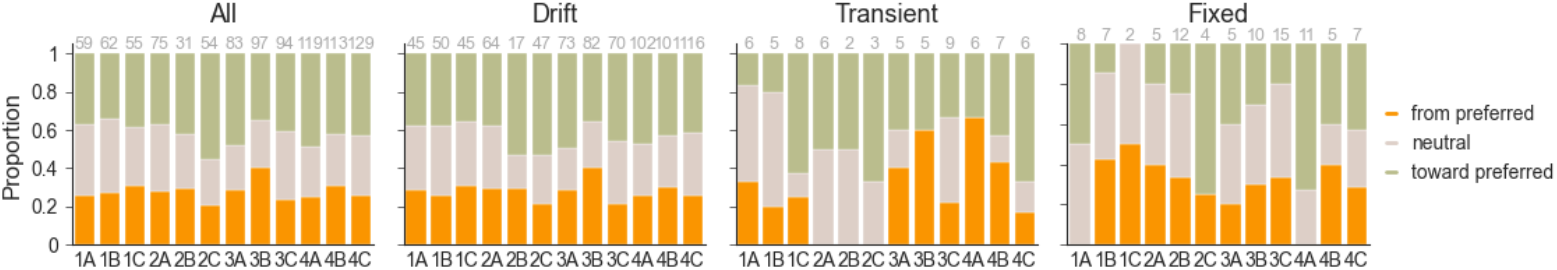
Direction of codon change and associated nucleotide change for synonymous SNVs. Direction of codon change relative to the preferred codon in the starting virus sequence for all and drift, transient, and fixed synonymous SNV per population.

**Extended Data Figure 7.**
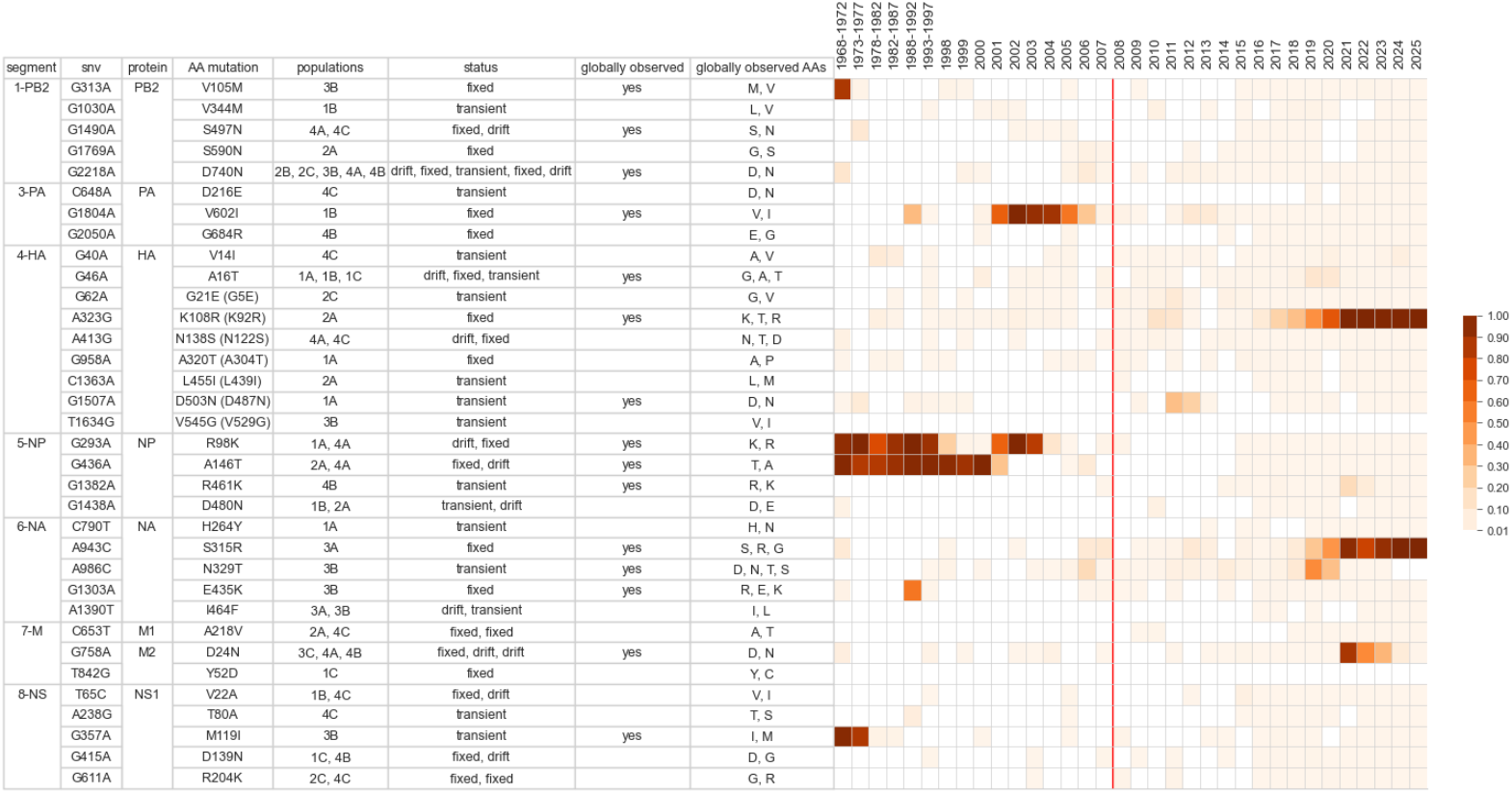
Overview of observed non-drift-like AA mutations in positions that are globally not conserved. Heatmap shows the global frequency of the observed AA mutation (e.g. for mutation V105M heatmap shows the global frequency of M). Global observation is defined as a frequency >10% in one the time windows. White cells indicate frequency <0.01. Red lines indicate 2008. Drift observation in populations have also been included if AA observation in another population was classified as non-drift-like. Status order matches the order of the populations. For HA substitution are labelled by sequential number with H3-numbering in brackets where possible.

**Extended Data Table 1.** Observed and simulated site-level parallelism for non-drift-like non-synonymous SNVs across genes. For each gene parallelism was assessed using two statistics: a site-count statistic, which counts the number of sites mutated in two or more populations; and a multiplicity-weighted statistic, which sums the number of populations sharing a mutated site across all parallel sites. Empirical *P*-values were derived from 10,000 permutations

| Segment | Gene | Observed<br>single sites | Median<br>simulated<br>single sites | 95% CI<br>simulated<br>single sites | Site-count |  |  |  | Multiplicity-weighted |  |  |  |
| --- | --- | --- | --- | --- | --- | --- | --- | --- | --- | --- | --- | --- |
|  |  |  |  |  | Observed<br>parallel<br>sites | Median<br>simulated<br>parallel sites | 95% CI<br>simulated<br>parallel sites | *P* | Observed<br>parallel<br>sites | Median<br>simulated<br>parallel sites | 95% CI<br>simulated<br>parallel sites | *P* |
| 1-PB2 | PB2 | 12 | 15 | 13 – 15 | 1 | 0 | 0 – 1 | 0.0560 | 3 | 0 | 0 – 2 | 0.0010 |
| 2-PB1 | PB1 | 10 | 10 | 8 – 10 | 0 | 0 | 0 – 1 | 1 | 0 | 0 | 0 – 2 | 1 |
| 3-PA | PA | 19 | 19 | 17 – 19 | 0 | 0 | 0 – 1 | 1 | 0 | 0 | 0 – 2 | 1 |
| 4-HA | HA | 12 | 14 | 12 – 14 | 1 | 0 | 0 – 1 | 0.0624 | 2 | 0 | 0 – 2 | 0.0624 |
| 5-NP | NP | 10 | 10 | 8 – 10 | 0 | 0 | 0 – 1 | 1 | 0 | 0 | 0 – 2 | 1 |
| 6-NA | NA | 17 | 17 | 15 – 17 | 0 | 0 | 0 – 1 | 1 | 0 | 0 | 0 – 2 | 1 |
| 7-M | M1 | 3 | 5 | 5 – 5 | 1 | 0 | 0 – 0 | 0.0183 | 2 | 0 | 0 – 0 | 0.0183 |
|  | M2 | 3 | 3 | 3 – 3 | 0 | 0 | 0 – 0 | 1 | 0 | 0 | 0 – 0 | 1 |
| 8-NS | NS1 | 8 | 10 | 8 – 10 | 1 | 0 | 0 – 1 | 0.0771 | 2 | 0 | 0 – 2 | 0.0771 |
|  | NEP | 1 | 3 | 3 – 3 | 1 | 0 | 0 – 0 | 0.0107 | 2 | 0 | 0 – 0 | 0.0107 |

**Extended Data Table 2.** Observed and simulated site-level parallelism for non-drift-like synonymous SNVs across genes. For each gene parallelism was assessed using two statistics: a site count statistic, which counts the number of sites mutated in two or more populations; and a multiplicity-weighted statistic, which sums the number of populations sharing a mutated site across all parallel sites. Empirical *P*-values were derived from 10,000 permutations.

| Segment | Gene | Observed<br>single sites | Median<br>simulated<br>single site | 95% CI<br>simulated<br>single sites | Count |  |  |  | Weighted |  |  |  |
| --- | --- | --- | --- | --- | --- | --- | --- | --- | --- | --- | --- | --- |
|  |  |  |  |  | Observed<br>multiple<br>sites | Median<br>simulated<br>multiple sites | 95% CI<br>simulated<br>multiple sites | *P* | Observed<br>multiple<br>sites | Median<br>simulated<br>multiple sites | 95% CI<br>simulated<br>multiple sites | *P* |
| 1-PB2 | PB2 | 21 | 23 | 19 – 23 | 1 | 0 | 0 – 2 | 0.3014 | 2 | 0 | 0 – 4 | 0.3014 |
| 2-PB1 | PB1 | 15 | 17 | 15 – 17 | 1 | 0 | 0 – 1 | 0.1854 | 2 | 0 | 0 – 2 | 0.1854 |
| 3-PA | PA | 26 | 28 | 24 – 28 | 1 | 0 | 0 – 2 | 0.4374 | 2 | 0 | 0 – 4 | 0.4374 |
| 4-HA | HA | 20 | 22 | 18 – 22 | 1 | 0 | 0 – 2 | 0.3703 | 2 | 0 | 0 – 4 | 0.3703 |
| 5-NP | NP | 20 | 22 | 18 – 22 | 1 | 0 | 0 – 2 | 0.4056 | 2 | 0 | 0 – 4 | 0.4056 |
| 6-NA | NA | 12 | 14 | 12 – 14 | 1 | 0 | 0 – 1 | 0.1845 | 2 | 0 | 0 – 2 | 0.1845 |
| 7-M | M1 | 10 | 12 | 9 – 12 | 1 | 0 | 0 – 1 | 0.2548 | 2 | 0 | 0 – 3 | 0.2548 |
|  | M2 | 5 | 5 | 3 – 5 | 0 | 0 | 0 – 1 | 1 | 0 | 0 | 0 – 2 | 1 |
| 8-NS | NS1 | 7 | 9 | 7 – 9 | 1 | 0 | 0 – 1 | 0.1490 | 2 | 0 | 0 – 2 | 0.1490 |
|  | NEP | 1 | 1 | 1 – 1 | 0 | 0 | 0 – 0 | 1 | 0 | 0 | 0 – 0 | 1 |

**Extended Data Table 3.** Nasal Human Airway Epithelial (HAE) donor information.

| Donor | Batch Number | Age | Sex | Origin | Smoker | Pathology | Month of cryo-preservation |
| --- | --- | --- | --- | --- | --- | --- | --- |
| 1 | 02AB038901 | 73 | M | White | No | No | January 2012 |
| 2 | AB047301 | 63 | M | White | No | No | October 2013 |
| 3 | AB060801 | 50 | M | White | No | No | November 2014 |
| 4 | AB063001 | 41 | F | White | No | No | February 2015 |

## Notes

https://github.com/AMC-LAEB/LTEE-flu

https://www.ncbi.nlm.nih.gov/bioproject/1549747

https://doi.org/10.5281/zenodo.23661876

### Competing Interest Statement

The authors have declared no competing interest.

