## Supplemental for "Long-term experimental evolution of seasonal influenza A/H3N2 virus reveals diverse evolutionary potential"

### Supplemental methods

#### 1. Quantification statistical support for identified candidate variants

To distinguish candidate SNVs (especially those present at minor frequencies) from sequencing errors, we evaluated an adapted version of the statistical framework described by Good, et al. (2017)[1]. This statistical framework utilises the temporal structure of our data, and evaluates the significance of each candidate SNV by comparing it to a null model where the reads supporting the SNVs are the result of random chance.

For each mutation $m$ in each population $p$, the alternate allele count and total sequencing depth at timepoint $t$ are defined as $A_{pmt}$ and $D_{pmt}$, respectively. In the null model, error probabilities are permitted to vary across samples and populations and are approximated using the pooled empirical allele frequencies, with a polarization correction applied to ensure agreement with the ancestral state. This polarization condition is defined as:

$$c = min\left\{1, \frac{1}{2\bar{f}}\right\}$$

where $\bar{f}_{pm} = \frac{\sum_t A_{pmt}}{\sum_t D_{pmt}}$. The null model error probabilities can then be generated via:

$$\hat{p}_{pmt} = cf_{pm\hat{\sigma}_p(t)}$$

Where $\hat{\sigma}(t)$ denotes a random permutation of the timepoints; removing all temporal information and thereby any genuine frequency dynamics while preserving the observed variation in $cf_{pmt}$. The alternate allele count under this error model is given by:

$$\hat{A}_{pmt} \sim \text{Binomial}(D_{pmt}, \hat{p}_{pmt})$$

We then quantified whether an observed SNV trajectory deviates from the null model using a composite test statistic that aggregates evidence across three features expected in true mutations. The first test statistic is (modified) autocorrelation [1] $C^*$, which captures the temporal correlation between closely sampled consecutive timepoints and is defined as:

$$C^* = \frac{\sum_t \sqrt{D_{t+1}D_t}(f_{t+1} - \bar{f})\theta(|A_{t+1} - \bar{f}D_{t+1}| - 1)\theta(|A_t - \bar{f}D_t| - 1)}{\bar{f}^2 \sum_t \sqrt{D_{t+1}D_t}}$$

where $\bar{f} = min\left\{\frac{\sum_t A_{pmt}}{\sum_t D_{pmt}}, \frac{1}{2}\right\}$ is average frequency weighted by sequencing depth, capped at 0.5 to match the polarization condition. $\theta(\cdot)$ is a Heaviside step function that ensures deviations from the average frequency $(f - \bar{f})$ are at least 1 read, reducing rounding artifacts.

The second test statistic is the derived allele sojourn weight[1] $I$ which captures the duration of the sojourn path of a true mutation; i.e., a true mutation will emerge and persist for some time after which it will either fix or go extinct. We identified all contiguous observations of two or more timepoints, $t = t_1, \ldots, t_2$ for which $f_t$ was larger than the threshold frequency $f^*$. This threshold frequency was set dynamically for each mutation via:

$$f^* = \begin{cases} \dfrac{\bar{f}}{1 + e^{+(n_0 - 0.3n)/5}} & if\ n_0 > 0.3n \\ \dfrac{\bar{f}}{1 + e^{-(n_0 - 0.3n)/5}} & if\ n_0 < 0.3n \end{cases}$$

Where $n$ and $n_0$ denote the total number of timepoints and the number of timepoints for which $A_{pmt} = 0$, respectively, and $\bar{f}$ is the capped average frequency. We then identified the observation with the largest derived allele sojourn weight:

$$I = \sum_{t=t_1}^{t_2} f_t - f^*$$

Finally, the average frequency relaxation time[1] $T$ captures the pattern of initial emergence of a true mutation. Here the assumption is that a true mutation starts with a zero or near-zero frequency and only later rises to higher frequency. Therefore, the average frequency of the first few timepoints should be lower than the average frequency over the entire time frame. For our data, we defined the relaxation time as:

$$T = max\left\{T : \frac{\sum_{t \le t'} A_{pmt}}{\sum_{t \le t'} D_{pmt}} \le 0.6 \cdot \frac{\sum_t A_{pmt}}{\sum_t D_{pmt}}\ \forall\, 3 \le t' < T\right\}$$

We set T = 0 if $n_0 > 0.3n$.

For each individual test statistic $T_k$, we calculated a single-statistic *P*-value:

$$P_k = \Pr\left[T_k\left(\{\hat{A}_{pmt}, D_{pmt}\}\right) \ge T_k\left(\{A_{pmt}, D_{pmt}\}\right)\right]$$

by simulating 10000 draws from $\{\hat{A}_{pmt}, D_{pmt}\}$. We then combined these individual *P*-values in a composite statistic[1]:

$$T\left(\{A_{pmt}, D_{pmt}\}\right) = \sum_k \theta\,(P^* - P_k) \log\left(\frac{1}{P_k}\right)$$

where $\theta(\cdot)$ is a Heaviside step function, and $P^* = (0.05)^{\frac{1}{3}}$, which ensure that statistic is dominated by largest deviations in one of the statistics, rather than smaller or moderate deviations in all of them. For this composite statistic we then calculated the composite *P*-value by again simulating 10000 draws from $\{\hat{A}_{pmt}, D_{pmt}\}$:

$$P_{pm} = \Pr\left[T\left(\{\hat{A}_{pmt}, D_{pmt}\}\right) \ge T_k\left(\{A_{pmt}, D_{pmt}\}\right)\right]$$

To obtain the final list of SNV, we performed multiple hypothesis correction on all trajectories where $P < 1$ via the Benjamini-Hochberg (BH) procedure[2] and with a false discovery rate (FDR) of 0.2.

#### 2. Successful infecting fraction estimation

To experimentally determine the fraction of infectious units that are successful in infecting the initial target cell population for the within-host simulations, we harvested and quantified the amount of unbound virus present in the wash following a 2-hour incubation at 33°C and 5% $CO_2$. In short, differentiated nasal HAE cells were inoculated with influenza A virus by adding 100µL of $10^4$/mL $TCID_{50}$ in HBSS to the apical side of the trans-well insert. Following inoculation, cells were incubated at 33°C and 5% $CO_2$ for 2 hours, after which the inoculum was removed and sampled, then the apical side was washed twice with HBSS which was also sampled. The viral titer of the unbound inoculum and wash samples was determined as described in the main text.

### Supplemental tables

**Table S1 Overview of samples per sequence batch and sequence platform samples were sequenced with**. For each sample, sequencing was performed for two technical replicates. * Indicates samples with low sequencing quality that were resequenced in a later batch.

| batch | Population | Passages | Sequencing platform |
|---|---|---|---|
| 1 | Starting | 0* | Illumina NovaSeq 6000 |
| 2 | 1A | 10, 20, 30, 40, 50, 60 ,70 | Illumina NovaSeq 6000 |
| | 1B | 10, 20, 30*, 40*, 50*, 60*, 70* | |
| | 1C | 10, 20, 30*, 40*, 50*, 60*, 70* | |
| | 2A | 10, 20, 30, 40*, 50, 60 ,70 | |
| | 2B | 10, 20, 30, 40, 50, 60 ,70 | |
| | 2C | 10, 20, 30, 40, 50, 60 ,70 | |
| | 3A | 10, 20, 30, 40, 50, 60 ,70 | |
| | 3B | 10, 20, 30, 40, 50, 60 ,70 | |
| | 3C | 10, 20, 30, 40, 50, 60 ,70 | |
| | 4A | 10, 20, 30, 40, 50, 60, 70* | |
| | 4B | 10, 20, 30, 40, 50, 60 ,70 | |
| | 4C | 10, 20, 30, 40, 50, 60 ,70 | |
| 3 | Starting | 0 | Illumina NovaSeq 6000 |
| | 2A | 30 | |
| | 4A | 70 | |
| 4 | 1A | 80 | Illumina NovaSeq X Plus |
| | 1B | 80 | |
| | 1C | 80 | |
| | 2A | 80 | |
| | 2B | 80 | |
| | 2C | 80 | |
| | 3A | 80 | |
| | 3B | 80 | |
| | 3C | 80 | |
| | 4A | 80 | |
| | 4B | 80 | |
| | 4C | 80 | |
| 5 | 1A | 90 | Illumina NovaSeq X Plus |
| | 1B | 90 | |
| | 1C | 90 | |
| | 2A | 90 | |
| | 2B | 90 | |
| | 2C | 90 | |
| | 3A | 90 | |
| | 3B | 90 | |
| | 3C | 90 | |
| | 4A | 90 | |
| | 4B | 90 | |
| | 4C | 90 | |
| 6 | 1A | 100 | Illumina NovaSeq X Plus |
| | 1B | 100 | |
| | 1C | 100 | |
| | 2A | 100 | |
| | 2B | 100 | |
| | 2C | 100 | |
| | 3A | 100 | |
| | 3B | 100 | |
| | 3C | 100 | |
| | 4A | 100 | |
| | 4B | 100 | |
| | 4C | 100 | |
| 7 | 1B | 30, 40, 50, 60, 70 | Illumina NovaSeq X Plus |
| | 1C | 30, 40, 50, 60, 70 | |

**Table S2 Parameters used in the within-host model.**

| Parameter | Meaning | Value (units) | Source |
|---|---|---|---|
| - | Number of hours per replicative generation | 6 hr | Baccam, et al. 2006[3] |
| - | Number of hours per passage | 72 hr | - |
| $C_0$ | Initial target cell population | $2 \cdot 10^5$ cells | Assumption |
| $V_{input}$ | Number of infectious units with which infection is started | 1000 virions | - |
| $f_{si}$ | Fraction of virions that are successful infecting the initial target cell population | 90% | Assumption, based on experiment validation (supplemental methods S2 and Table S10) |
| $r$ | Number of virions produced by an infected cell | 100 | Frensing, et al. 2016[4] |
| $R_0$ | Within-host basic reproduction number | 5 | Hadjichrysanthou, et al. 2016[5] |
| $d$ | Per capita decay rate | 2 per generation | Han, et al. 2021[6] |
| $L$ | Binary genome length | cds of genome segments | - |
| $\mu$ | Per-site per generation mutation rate | $3 \cdot 10^{-6}$ per site per generation | Assumption estimated from data |
| $f_{apical}$ | Fraction of virions that are release into the apical compartment of the transwell plate | 50% | Assumption |
| $f_{wash}$ | Fraction of virions that are recovered by performing apical washes | 50% | Assumption |

**Table S3 Number of non-drift-like non-synonymous SNVs acquired in each genome segment per experimental population.**

| Population | 1-PB2 | 2-PB1 | 3-PA | 4-HA | 5-NP | 6-NA | 7-M | 8-NS |
|---|---|---|---|---|---|---|---|---|
| 1A | 0 | 1 | 1 | 4 | 0 | 3. | 0 | 0 |
| 1B | 2 | 1 | 2 | 1 | 3 | 1 | 0 | 1 |
| 1C | 1 | 2 | 2 | 2 | 0 | 0 | 2 | 2 |
| 2A | 1 | 0 | 1 | 2 | 2 | 1 | 1 | 1 |
| 2B | 1 | 1 | 1 | 1 | 1 | 0 | 0 | 0 |
| 2C | 2 | 0 | 3 | 1 | 0 | 1 | 1 | 2 |
| 3A | 0 | 0 | 2 | 0 | 0 | 2 | 0 | 1 |
| 3B | 2 | 0 | 3 | 1 | 0 | 5 | 0 | 1 |
| 3C | 1 | 3 | 4 | 0 | 1 | 2 | 2 | 1 |
| 4A | 3 | 1 | 2 | 0 | 2 | 0 | 1 | 0 |
| 4B | 0 | 1 | 3 | 0 | 1 | 1 | 0 | 0 |
| 4C | 2 | 0 | 2 | 2 | 0 | 1 | 1 | 2 |

**Table S4 Number of non-drift-like synonymous SNVs acquired in each genome segment per experimental population**.

| Population | 1-PB2 | 2-PB1 | 3-PA | 4-HA | 5-NP | 6-NA | 7-M | 8-NS |
|---|---|---|---|---|---|---|---|---|
| 1A | 2 | 0 | 2 | 2 | 3 | 0 | 2 | 1 |
| 1B | 1 | 2 | 2 | 1 | 3 | 0 | 1 | 2 |
| 1C | 1 | 3 | 0 | 3 | 1 | 0 | 2 | 0 |
| 2A | 0 | 1 | 3 | 0 | 1 | 3 | 1 | 0 |
| 2B | 4 | 1 | 2 | 2 | 3 | 1 | 1 | 0 |
| 2C | 0 | 1 | 4 | 1 | 1 | 0 | 0 | 0 |
| 3A | 0 | 2 | 1 | 1 | 2 | 0 | 2 | 2 |
| 3B | 4 | 1 | 2 | 2 | 1 | 1 | 1 | 1 |
| 3C | 2 | 1 | 6 | 4 | 3 | 4 | 2 | 1 |
| 4A | 2 | 2 | 2 | 2 | 2 | 2 | 4 | 1 |
| 4B | 4 | 2 | 0 | 2 | 1 | 1 | 1 | 1 |
| 4C | 3 | 1 | 4 | 2 | 1 | 2 | 0 | 0 |

**Table S5 Number of transitions and transversion substitution and the resulting estimated transition/transversion (Ti/Tv) ratio per population, as well as the P-value support from comparison with the expected Ti/Tv ratio**.

| population | # Transitions | # Transversions | # total substitutions | Estimated Ti/Tv ratio | P (two-side binomial tests) |
|---|---|---|---|---|---|
| 1A | 37 | 4 | 41 | 9.25 | P<0.05 |
| 1B | 44 | 1 | 45 | 44 | P<0.001 |
| 1C | 40 | 5 | 45 | 8 | P<0.05 |
| 2A | 55 | 3 | 58 | 18.33 | P<0.001 |
| 2B | 13 | 3 | 16 | 4.33 | 0.77 |
| 2C | 40 | 2 | 42 | 20 | P<0.01 |
| 3A | 63 | 4 | 67 | 15.75 | P<0.001 |
| 3B | 68 | 7 | 75 | 9.71 | P<0.01 |
| 3C | 60 | 4 | 64 | 15 | P<0.001 |
| 4A | 85 | 8 | 93 | 10.63 | P<0.001 |
| 4B | 82 | 11 | 93 | 7.45 | P<0.01 |
| 4C | 103 | 6 | 109 | 17.167 | P<0.001 |

**Table S6 Shared non-drift-like non-synonymous SNVs**. SNVs classified as non-drift-like (i.e. transient or fixed) in at least two experimental population are included in the table. Drift classifications of the included SNVS are also reported in the table.

| Segment | SNV | Protein | AA mutation | Population | Classification | Hitchhiker | Cooccurrence | Mutation bias |
|---|---|---|---|---|---|---|---|---|
| 1-PB2 | G2218A | PB2 | D740N | 2B | drift | | | G>A |
| | | | | 2C | fixed | | high confidence | |
| | | | | 3B | transient | | | |
| | | | | 4A | fixed | | | |
| | | | | 4B | drift | | | |
| 4-HA | G46A | HA | A16T | 1A | drift | | | G>A |
| | | | | 1B | fixed | | | |
| | | | | 1C | transient | | | |
| 7-M | C653T | M1 | A218V | 2A | fixed | | | C>T |
| | | | | 4C | fixed | | | |
| 8-NS | G611A | NS1 | R204K | 2C | fixed | | | G>A |
| | | | | 4C | fixed | | | |
| | | NEP | E52K | 4C | fixed | | | |
| | | | | 2C | fixed | | | |

**Table S7 Shared non-drift-like synonymous SNVs**. SNVs classified as non-drift-like (i.e. transient or fixed) in at least two experimental population are included in the table. Drift classifications of the included SNVS are also reported in the table.

| Segment | SNV | Protein | AA mutation | Population | Classification | Hitchhiker | Co-occurrence | Mutation bias |
|---|---|---|---|---|---|---|---|---|
| 1-PB2 | C1971T | PB2 | N657N | 4A | fixed | high confidence | potential | C>T |
| | | | | 4B | transient | | | C>T |
| 2-PB1 | C1243T | PB1 | L415L | 1C | fixed | high confidence | | C>T |
| | | | | 3A | fixed | | | |
| | | | | 4A | drift | | | |
| 4-HA | G1290A | HA | E430E (H3: E414E) | 1C | transient | | | G>A |
| | | | | 3A | drift | | | |
| | | | | 3C | transient | | | |
| 5-NP | G450A | NP | R150R | 3A | fixed | | | G>A |
| | | | | 4A | transient | | | |
| | | | | 4B | drift | | | |
| 6-NA | C1389T | NA | D463D | 1B | drift | | | C>T |
| | | | | 3C | fixed | potential | potential | |
| | | | | 4A | fixed | | | |
| | | | | 4B | drift | | | |
| 7-M | G324A | M1 | T108T | 1A | drift | | | G>A |
| | | | | 1C | transient | | | |
| | | | | 3B | transient | | | |
| | | | | 4A | drift | | | |
| | | | | 4B | drift | | | |
| 8-NS | A300G | NS1 | R100R | 1B | fixed | | potential | |
| | | | | 4A | transient | | | |

**Table S8 Global conservation and number of unique AA mutations observed within and outside of globally conserved sites per gene**.

| Gene | Global conservation | Number of unique AA mutations observed in globally conserved sites | Number of unique AA mutations observed in globally not conserved sites |
|---|---|---|---|
| PB2 | 91.97 % (699/760) | 8/13 (61.5%) | 5/13 (38.5%) |
| PB1 | 94.3 % (715/758) | 10/10 (100%) | 0/10 (0 %) |
| PA | 92.2 % (661/717) | 16/19 (84.2 %) | 3/19 (15.8%) |
| HA | 74.4 % (422/567) | 4/13 (30.8%) | 9/13 (69.2%) |
| NP | 89.2 % (445/499) | 6/10 (60.0%) | 4/10 (40.0%) |
| NA | 75.5 % (355/470) | 12/17 (70.6%) | 5/17 (29.4%) |
| M1 | 92.9 % (235/253) | 3/4 (75 %) | 1/4 (25 %) |
| M2 | 76.5 % (75/98) | 1/3 (33.3%) | 2/3 (66.7%) |
| NS1 | 70.2 % (167/238) | 4/9 (44.4%) | 5/9 (55.6%) |
| NEP | 92.1 % (117/127) | 2/2 (100%) | 0/2 (0 %) |

**Table S9 Overview of observed non-drift-like AA mutations in globally conserved positions per gene**. Drift observation in populations have also been included if AA observation in another population was classified as non-drift-like.

| Segment | SNV | Protein | AA mutation | Populations | Status | Globally observed AAs |
|---|---|---|---|---|---|---|
| 1-PB2 | C128T | PB2 | P43L | 1C | fixed | P |
| | A526G | | I176V | 3C | transient | I |
| | A874G | | T292A | 2C | fixed | T |
| | G1476T | | E492D | 4A | fixed | E |
| | G1750A | | V584I | 4C | transient | V |
| | A1846G | | I616V | 2B | fixed | I |
| | G1864A | | A622T | 1B | fixed | A |
| | G2149A | | A717T | 4C | transient | A |
| 2-PB1 | A543G | PB1 | I181M | 2B | fixed | I |
| | T639A | | N213K | 4A | fixed | N |
| | A1015G | | I339V | 3C | fixed | I |
| | G1196A | | G399D | 1A, 4C | transient, drift | G |
| | C1360T | | P454S | 1A, 4B | drift, transient | P |
| | G1569A | | M523I | 1C | fixed | M |
| | T1919C | | V640A | 3C | transient | V |
| | G2117A | | R706K | 3C | transient | R |
| | C2165G | | A722G | 1B | transient | A |
| | T2188A | | F730I | 1C | transient | F |
| 3-PA | A112G | PA | I38V | 4B | fixed | I |
| | G258A | | M86I | 4A | transient | M |
| | G465A | | M155I | 2A, 4A | drift, fixed | M |
| | C530T | | T177I | 3A | fixed | T |
| | G575A | | R192H | 2C | fixed | R |
| | T968C | | V323A | 3B, 4C | fixed, drift | V |
| | C1207A | | L403I | 1B | transient | L |
| | G1255A | | D419N | 2C | fixed | D |
| | G1380A | | M460I | 3B | transient | M |
| | C1434A | | D478E | 3C | fixed | D |
| | G1561T | | V521L | 4B | fixed | V |
| | C1601T | | P534L | 3A | fixed | P |
| | G1785A | | M595I | 1C, 3B | fixed, drift | M |
| | A1817G | | D606G | 2B | fixed | D |
| | G1880A | | G627E | 1C | fixed | G |
| | T1894C | | S632P | 1A | transient | S |
| 4-HA | G106A | HA | V20I (V36I) | 1B, 2B | drift, fixed | V |
| | T947C | | I300T (I316T) | 1C | fixed | I |
| | A1242T | | E398D (E414D) | 1A | fixed | E |
| | A1300C | | I418L (I434L) | 1A | transient | I |
| 5-NP | G541A | NP | A181T | 2A | fixed | A |
| | T1112C | | M371T | 1B | transient | M |
| | G1113A | | M371I | 1B | transient | M |
| | T1442C | | M481T | 4A | fixed | M |
| | G1477A | | A493T | 2B, 4B, 4C | fixed, drift, drift | A |
| | G1489T | | D497Y | 3C | transient | D |
| 6-NA | C295T | NA | P99S | 1A | transient | P |
| | G373A | | D125N | 2A, 4B | fixed, drift | D |
| | G529T | | A177S | 1B | transient | A |
| | G574A | | V192I | 3A | transient | V |
| | C584T | | T195M | 3C | transient | T |
| | A784G | | I262V | 3B | transient | I |
| | T935C | | I312T | 4C | transient | I |
| | G1070A | | G357E | 3B | fixed | G |
| | G1086A | | M362I | 1C, 3C | drift, fixed | M |
| | G1186A | | V396I | 4B | transient | V |
| | A1327G | | I443V | 1A | fixed | I |
| | G1342A | | G448S | 2C | fixed | G |
| 7-M | C328T | M1 | H110Y | 2C | fixed | H |
| | G412A | | V138I | 3C | fixed | V |
| | G544A | | A182T | 4A | fixed | A |
| | G773A | M2 | A29T | 1C, 4B | transient, drift | A |
| 8-NS | G226A | NS1 | A76T | 3A | fixed | A |
| | G256A | | A86T | 1C, 2C | transient, drift | A |
| | G299A | | R100K | 2A, 3C | drift, fixed | R |
| | G406A | | V136M | 2C | fixed | V |
| | G611A | NEP | E52K | 2C, 4C | fixed, fixed | E |
| | A815G | | T120A | 2A | transient | T |

**Table S10 Estimated successful infecting fractions**.

| HAE Donor | Technical Replicate | Log $TCID_{50}$ titer/mL | | | Number of infectious units | | | Successful infection % |
|---|---|---|---|---|---|---|---|---|
| | | Input Inoculum | Unbound Inoculum after 2 hours | Wash | Input | Recovered from wash and harvest | Successful in infection | |
| 1 | 1 | 4 | undetected | undetected | 1000 | 0 | 1000 | 100 |
| 1 | 2 | 4 | 3.37 | undetected | 1000 | 234.4 | 765.6 | 76.5 |
| 2 | 1 | 4 | 2.64 | undetected | 1000 | 43.7 | 956.3 | 95.6 |
| 3 | 1 | 4 | undetected | undetected | 1000 | 0 | 1000 | 100 |
| 3 | 2 | 4 | 2.54 | undetected | 1000 | 34.7 | 965.3 | 96.5 |
| 4 | 1 | 4 | undetected | undetected | 1000 | 0 | 1000 | 100 |
| 4 | 2 | 4 | undetected | undetected | 1000 | 0 | 1000 | 100 |
| 4 | 3 | 4 | undetected | undetected | 1000 | 0 | 1000 | 100 |
| 4 | 4 | 4 | undetected | undetected | 1000 | 0 | 1000 | 100 |

**Table S11 Number of global surveillance sequences included in comparison per time window per segment**.

| Time window | 1-PB2 | 2-PB1 | 3-PA | 4-HA | 5-NP | 6-NA | 7-M | 8-NS |
|---|---|---|---|---|---|---|---|---|
| 1968-1972 | 138 | 139 | 140 | 188 | 149 | 149 | 151 | 154 |
| 1973-1977 | 58 | 56 | 56 | 104 | 62 | 62 | 60 | 59 |
| 1978-1982 | 38 | 38 | 36 | 97 | 37 | 42 | 41 | 40 |
| 1982-1987 | 56 | 56 | 50 | 213 | 52 | 59 | 56 | 59 |
| 1988-1992 | 122 | 120 | 122 | 378 | 128 | 135 | 128 | 129 |
| 1993-1997 | 350 | 346 | 344 | 849 | 360 | 431 | 416 | 354 |
| 1998 | 57 | 59 | 56 | 363 | 60 | 126 | 123 | 59 |
| 1999 | 139 | 140 | 139 | 514 | 141 | 215 | 220 | 141 |
| 2000 | 138 | 139 | 138 | 195 | 137 | 148 | 138 | 139 |
| 2001 | 56 | 55 | 52 | 114 | 53 | 75 | 59 | 55 |
| 2002 | 184 | 184 | 180 | 342 | 180 | 228 | 188 | 184 |
| 2003 | 309 | 309 | 298 | 463 | 308 | 398 | 320 | 313 |
| 2004 | 256 | 259 | 268 | 403 | 253 | 312 | 282 | 253 |
| 2005 | 261 | 260 | 259 | 376 | 257 | 310 | 296 | 262 |
| 2006 | 64 | 64 | 66 | 294 | 61 | 188 | 118 | 64 |
| 2007 | 231 | 236 | 226 | 791 | 231 | 486 | 464 | 238 |
| 2008 | 79 | 77 | 49 | 438 | 77 | 364 | 247 | 79 |
| 2009 | 373 | 370 | 360 | 1118 | 364 | 920 | 646 | 369 |
| 2010 | 62 | 63 | 63 | 190 | 63 | 139 | 121 | 58 |
| 2011 | 140 | 140 | 140 | 332 | 135 | 239 | 228 | 138 |
| 2012 | 245 | 245 | 174 | 573 | 211 | 305 | 305 | 245 |
| 2013 | 311 | 350 | 149 | 576 | 301 | 418 | 366 | 316 |
| 2014 | 412 | 444 | 449 | 1101 | 463 | 604 | 522 | 460 |
| 2015 | 1287 | 1303 | 1318 | 2427 | 1363 | 1553 | 1421 | 1352 |
| 2016 | 2377 | 2370 | 2415 | 3438 | 2435 | 2838 | 1744 | 2435 |
| 2017 | 5200 | 5178 | 5194 | 6920 | 2335 | 5997 | 5297 | 5225 |
| 2018 | 4260 | 4226 | 4249 | 5361 | 4292 | 4753 | 4323 | 4256 |
| 2019 | 7516 | 7432 | 7567 | 19055 | 7651 | 8762 | 7750 | 7557 |
| 2020 | 1061 | 1033 | 1129 | 1567 | 1070 | 1430 | 1094 | 1069 |
| 2021 | 4575 | 4413 | 4794 | 5454 | 4795 | 5166 | 4904 | 4838 |
| 2022 | 20065 | 19142 | 20423 | 25011 | 21025 | 23300 | 21920 | 21337 |
| 2023 | 7608 | 7312 | 7780 | 9395 | 8098 | 9047 | 8417 | 8099 |
| 2024 | 14737 | 14136 | 15314 | 17161 | 15630 | 16751 | 16172 | 15693 |
| 2025 | 29694 | 29044 | 30000 | 32856 | 31225 | 32201 | 31806 | 30886 |

#### Supplemental figures

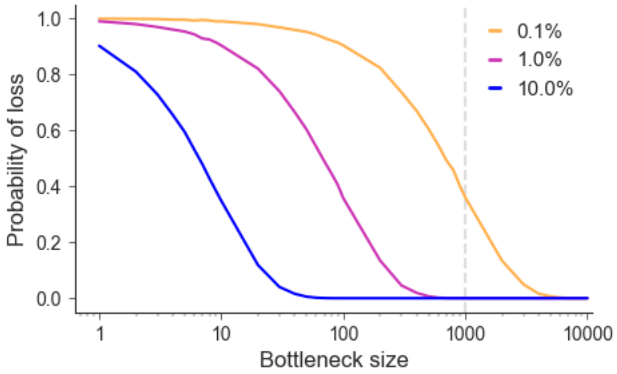


**Figure S1 Impact of passage bottleneck of loss of viral diversity**. Each coloured line represents a different variant frequency. Vertical dashed grey line represents the 1000 virion passage bottleneck maintained during the experiment.

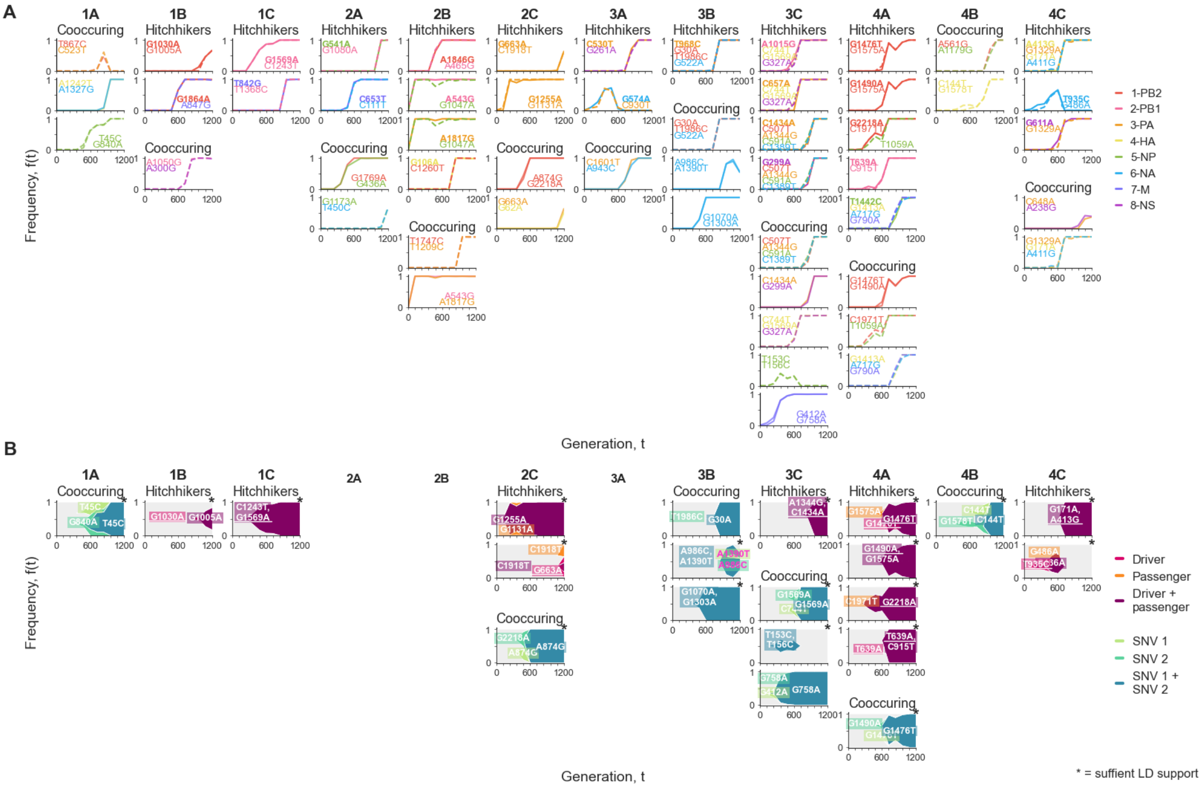


**Figure S2 Identified putative hitchhiker (driver-passenger(s) pair) and cooccurring amongst non-drift-like SNVs per population**. **A**. Frequency trajectories of identified putative hitchhiker and cooccurring SNVs. SNVs are coloured by the segment on which they occurred. Solid and dotted lines indicate non-synonymous and synonymous SNVs, respectively. **B**. Haplotype dynamics of the identified putative hitchhiker and cooccurring SNVs occurring on the same genome segments. Asterisk indicates sufficient support ($D' \geq 0.95$) from genetic linkage from linkage disequilibrium (LD) calculations. Pink annotation indicates lost SNV in haplotype.

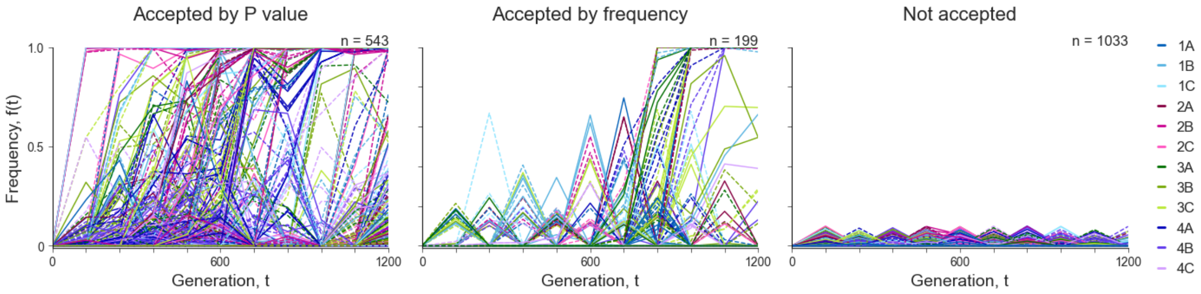


**Figure S3 The allele frequency trajectories of all *de novo* single nucleotide variants (SNVs) detected in the experimental populations separate by classified error model acceptance and/or frequency acceptance.** SNV trajectories accepted by frequency have a *P*-value above 0.05, yet their frequency reached above the acceptance threshold of 10%. SNV trajectories are coloured by populations.

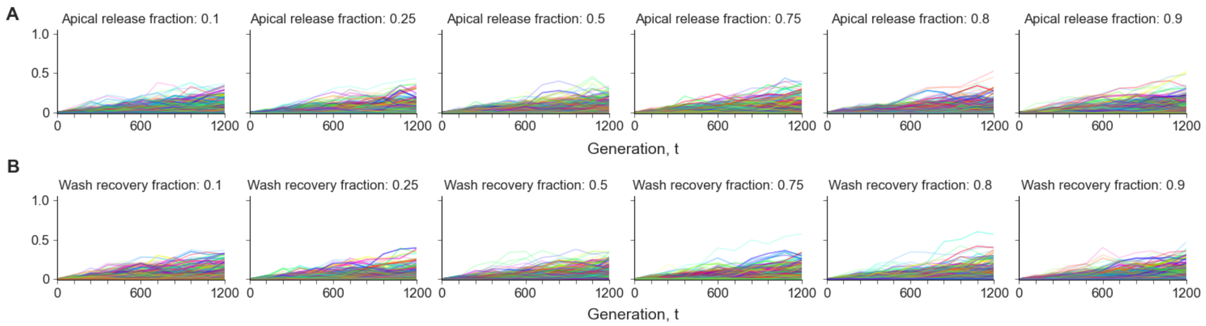


**Figure S4 Simulated frequency trajectories with varying apical release fractions (A) and wash recovery fractions (B).** For each simulation run we performed 10 simulations of 100 passages on a genome with 2,281 binary loci (max genome length simulated). All other parameter values were set to the default given in Supplemental Table S2, including wash recovery fraction in panel A and apical release fraction in panel B. SNV trajectories are coloured by simulation for visualization.

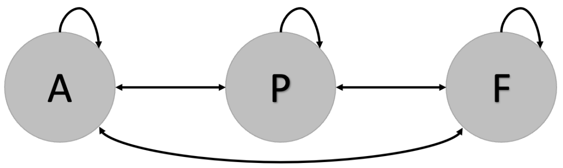


**Figure S5 Schematic of the SNV trajectory HMM.** Schematic shows possible transitions between the ancestral ***A***, polymorphic ***P***, and fixed ***F*** Markov states.

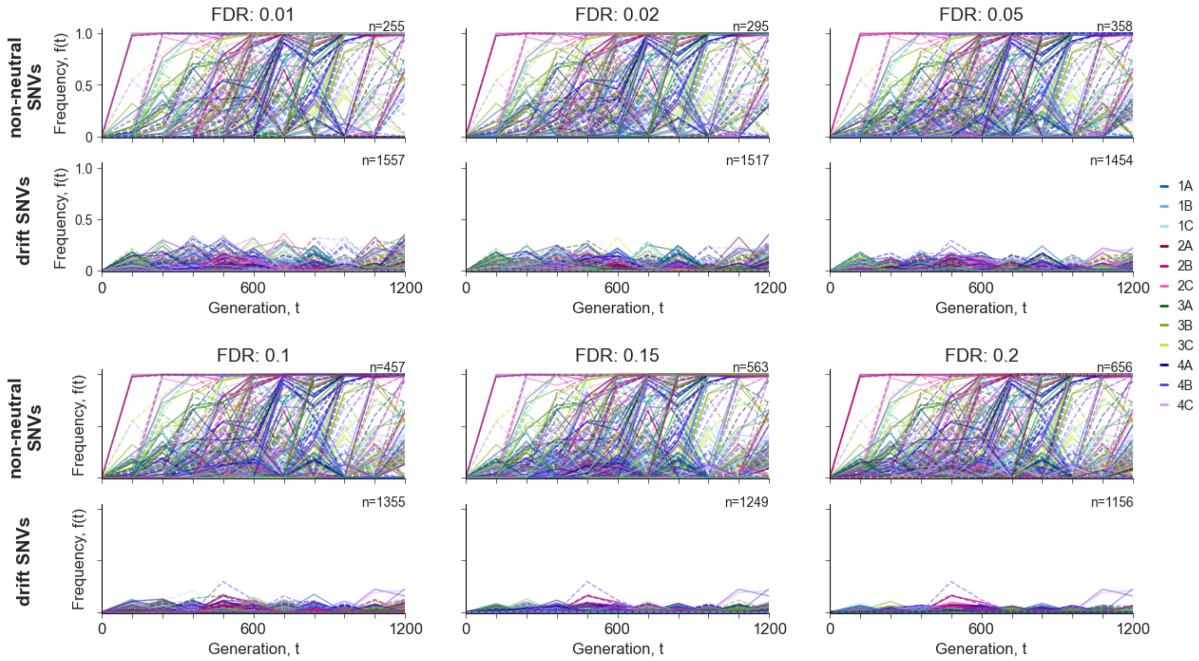


**Figure S6 Sensitivity analysis of false discovery rates (FDR) for the statistical identification of non-drift-like SNVs.** SNV trajectories are coloured by population.
